# Distinct colorectal cancer genotypes shape microbial ecosystems and reveal stage-specific microbiota dependencies

**DOI:** 10.64898/2026.07.06.736293

**Authors:** Valentina Brunner, Nicholas Bodenstein, Antonio E. Zaurito, Miguel G. Silva, Frederic Saab, Fabian Springer, Fabio Boniolo, Moritz Jesinghaus, Anantharamanan Rajamani, Aritra Mahapatra, Tanja Groll, Nicole A. Schmid, Rupert Öllinger, Chen Meng, Sabine Schwamberger, Julius C. Fischer, Georg Zeller, Karin Kleigrewe, Katja Steiger, Klaus Neuhaus, Dirk Haller, Roland Rad, Dieter Saur, Markus Tschurtschenthaler

## Abstract

The gut microbiota has emerged as an important modifier of colorectal cancer (CRC), yet how tumor genotype influences host-microbiota interactions and whether microbial signals are required throughout tumor progression remain unclear. Here, we combined genetically engineered mouse models, microbial multi-omics and a germ-free-compatible orthotopic transplantation system to define the functional contribution of the microbiota across distinct stages of CRC evolution. Across multiple CRC genotypes, we identified tumor-associated microbial ecosystem states characterized by distinct taxonomic, functional and metabolic configurations. To directly test their contribution to tumor progression, we established the first orthotopic CRC transplantation platform compatible with long-term experimentation in germ-free mice, enabling side-by-side comparison of genetically identical tumors in the presence or absence of microbiota. Using organoids spanning low-grade adenoma, high-grade adenoma and adenocarcinoma states, we found that the dependence on the presence of microbiota progressively decreases during malignant evolution. Whereas adenoma-derived organoids exhibited profound dependence on microbial exposure and failed or were markedly impaired in establishing tumors under germ-free conditions, adenocarcinoma organoids engrafted and metastasized in both germ-free and specific pathogen-free (SPF) hosts. Unexpectedly, comprehensive histological, immunological and transcriptomic analyses revealed highly similar tumor ecosystem states in advanced tumors arising under both microbial conditions, arguing against broad immune or epithelial defects as a primary explanation for the observed phenotype. Together, our findings demonstrate that distinct oncogenic drivers establish specific microbial ecosystem states and reveal a stage-dependent role of the microbiota during colorectal tumorigenesis. Whereas microbial signals are critical during early stages of tumor progression and may promote malignant transformation, advanced tumors progressively acquire microbiota-independent growth programs and increasingly impose genotype-specific ecological signatures on the surrounding microbial ecosystem. More broadly, we establish a versatile framework for the causal dissection of tumor-microbiota interactions in cancer.

## Introduction

Colorectal cancer (CRC) arises through multiple molecular routes characterized by distinct combinations of genetic alterations, histopathological features and clinical behaviors.^1,2^ Large-scale molecular classification efforts have established that CRC comprises biologically diverse disease entities that differ in oncogenic signaling, immune infiltration, stromal composition and therapeutic vulnerabilities. Classical adenoma-carcinoma progression is most commonly initiated by disruption of the APC/WNT pathway,^3^ whereas serrated neoplasia is frequently driven by activation of the MAPK and PI3K signaling axes through oncogenic *KRAS*, *BRAF* or *PIK3CA* mutations.^4–6^ These distinct oncogenic programs not only shape tumor cell-intrinsic phenotypes but also profoundly influence the surrounding tissue ecosystem.^7–10^

The intestinal microbiota has emerged as an important component of this ecosystem.^11,12^ Multiple studies have linked CRC to microbial dysbiosis, enrichment of specific bacterial taxa and altered microbial metabolite production. Experimental work has further demonstrated that individual microorganisms, microbial communities and microbially derived metabolites can promote intestinal tumorigenesis through effects on epithelial signaling, inflammation, DNA damage and immune regulation.^13–17^ Moreover, microbial composition has been associated with therapeutic response and clinical outcome in both patients and preclinical models.^18–20^ Collectively, these observations have led to the concept that CRC develops within a dynamic host-microbiota ecosystem in which tumor cells and microorganisms continuously influence one another.

Despite growing evidence linking the microbiota to CRC, several fundamental questions remain unresolved. First, it is unclear whether distinct oncogenic drivers instruct specific microbial ecosystem states during tumor evolution. Most studies have focused on microbial alterations associated with established tumors,^13,21–23^ making it difficult to distinguish whether microbial changes arise as a consequence of tumor growth or reflect genotype-specific ecological remodeling by the epithelium itself. Second, although numerous associations between microbial dysbiosis and CRC have been reported,^24–26^ the functional contribution of the microbiota across different stages of tumor progression remains poorly understood. Whether microbial signals are equally required throughout tumor evolution, or whether progressing tumors eventually acquire microbiota-independent growth programs, is unknown. Addressing these questions has been hindered by limitations of existing experimental systems. Germ-free mouse models provide a powerful approach for interrogating microbiota function but are technically challenging to combine with genetically engineered cancer models, which often require extensive breeding strategies and are difficult to maintain under germ-free conditions. Furthermore, most intestinal cancer models predominantly develop tumors in the small intestine rather than the colon,^27^ limiting their utility for studying tumor-microbiota interactions within the native colorectal microenvironment. Importantly, existing approaches generally do not permit direct side-by-side comparison of genetically identical tumors in the presence or absence of microbiota, making causal interrogation of microbiota dependence difficult.

Here, we combined genetically engineered mouse models representing major molecular routes of CRC with microbial multi-omics profiling and a germ-free-compatible orthotopic transplantation platform to define the relationship between tumor genotype, microbial ecosystem state and microbiota-dependent tumor progression. We show that distinct oncogenic drivers establish reproducible microbiota configurations prior to and during tumor development, generating genotype-specific microbial and metabolic ecosystem states. To determine the functional significance of these ecosystems, we developed the first orthotopic colorectal cancer transplantation system compatible with long-term experimentation in fully immunocompetent germ-free mice, enabling direct side-by-side comparison of genetically and molecularly defined tumors under germ-free and conventional conditions. Using this platform, we demonstrate that microbiota dependence is strongly stage-specific: whereas adenoma-derived tumors exhibit profound dependence on microbial exposure, advanced carcinomas progress largely independently of the microbiota. Together, our findings identify oncogenic genotype as a key determinant of CRC-associated microbial ecosystems and reveal that the requirement for microbial signals is progressively lost during malignant evolution.

## Results

### Oncogenic drivers establish distinct microbial ecosystem states during intestinal tumorigenesis

CRC is a very heterogeneous disease with different molecular routes and sets of genetic events, accompanied by alterations of the immune response and influences of exogenous factors, and represents a challenge for personalized therapeutic approaches.^1,28,29^ Over the past 10 years, many groups and consortia have made great efforts to stratify CRC into molecular subgroups, which is the key for identifying and developing subtype-specific therapeutic targets and responses to therapy.^1,2,29^ Today, three main routes of CRC development are currently recognised; i) the classical route that exhibits initial mutations in the tumor suppressor gene *Adenomatous Polyposis Coli* (*APC*), which lead to β-catenin accumulation in the nucleus and constitutive activation of the Wnt pathway;^30^ ii) the serrated pathway that exhibits activation of the RAS-RAF-MEK-ERK axis or PI3K/AKT pathway through expression of oncogenic *KRAS*^G12D^, *BRAF*^V600E^ or *PIK3CA*^H1047R^;^1,31,32^ and iii) the colitis-associated route that exhibits initial mutations in the tumor suppressor gene *TRP53*, followed by expression of oncogenic KRAS^G12D^ and finally mutations in *APC*.^33^ In order to study tumor-promoting cell-intrinsic and extrinsic mechanisms in subtypes of CRC, we and others generated mouse models for classical, serrated and mucinous intestinal cancer using a Cre recombinase that is under the intestinal epithelial cell (IEC)-specific promoter *Villin* (hereafter referred to as *V*-Cre) together with Cre-activatable oncogenic *Kras*^G12D^,^31^ *Braf*^V637E^ (the mouse orthologue of the human oncogenic BRAF p.V600E oncogenic mutation)^32^ and *Pik3ca*^H1047R^ ^34^ alleles **(Fig. 1a)**. Moreover, we also generated mice with IEC-specific loss of the tumor suppressor APC (*Apc*^fl/wt^).^35^ Of note, *Apc*^fl/wt^ mice exhibited a median survival time of 319 days (*n* = 120) and developed several tumor lesions via the classical adenoma-carcinoma route, while *Braf*^V637E^, *Kras*^G12D^ and *Pik3ca*^H1047R^ mice exhibited a median survival time of 551 days (*n* = 92), 643 days (*n* = 89) and 642 days (*n* = 117), respectively, and showed increased epithelial regeneration that spontaneously progressed to hyperplasia (preneoplastic lesions of the serrated route) and serrated adenoma, which could then further develop to serrated carcinoma **(Suppl. Fig. 1a-d)**. Tumors in all models are primarily located in the small intestine **(Suppl. Fig. 1e)**.

**Figure 1.**
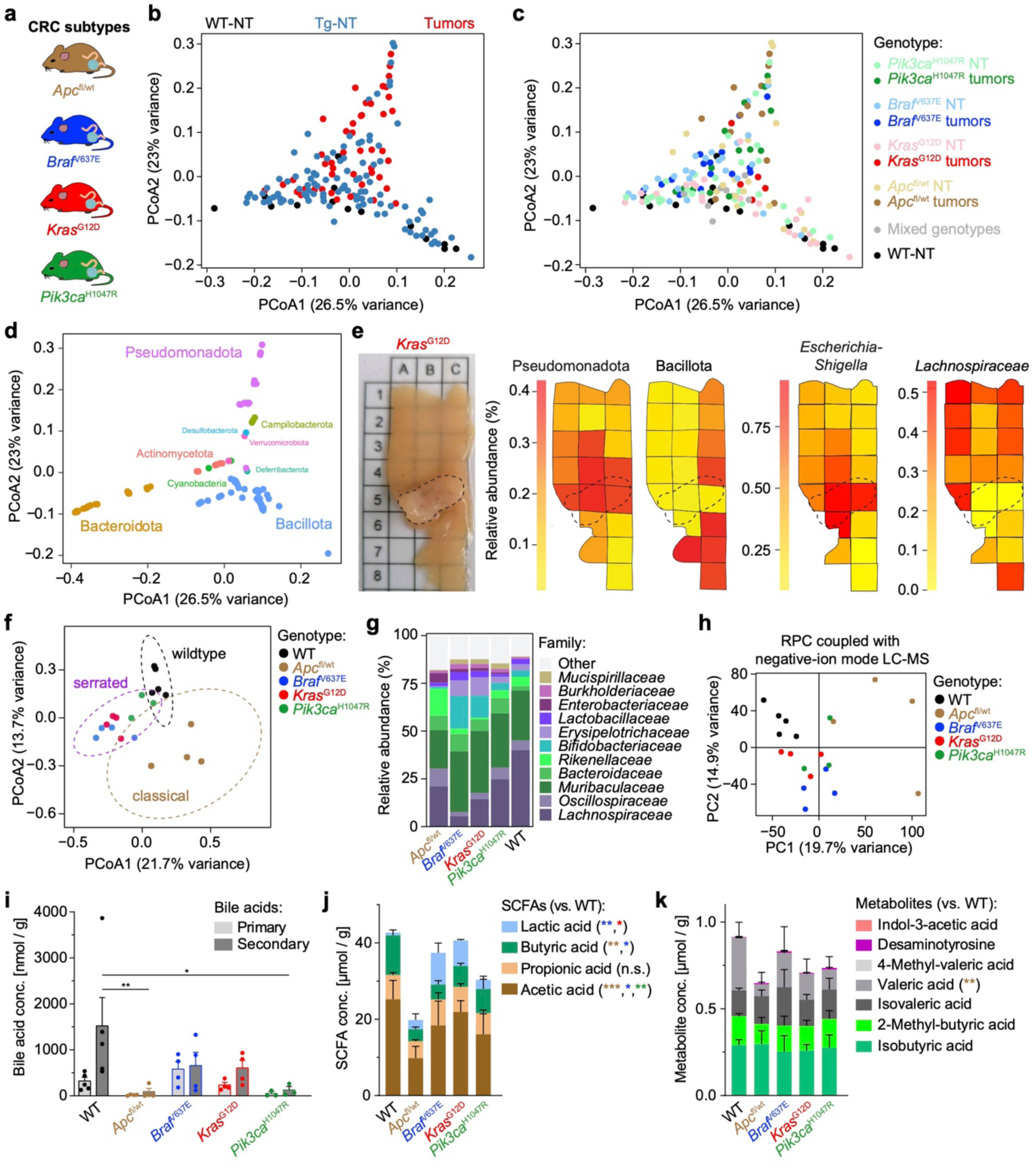
Distinct oncogenic drivers establish genotype-specific microbial ecosystem states during intestinal tumorigenesis. **(a)** Schematic overview of intestinal epithelial cell (IEC)-specific genetically engineered mouse models generated using *Villin*-Cre (V-Cre) representing the major molecular routes of colorectal tumorigenesis, including *Apc*^fl/wt^ (classical adenoma-carcinoma pathway) and *Braf*^V637E^, *Kras*^G12D^ and *Pik3ca*^H1047R^ (serrated pathway). **(b-d)** Principal coordinates analysis (PCoA) of mucosa-associated microbial communities from small intestinal samples of wild-type (WT) mice and non-tumorous (Tg-NT) and tumor tissues from transgenic mice representing the molecular CRC subtypes. Samples are shown according to tissue type and progression stage **(b)**, oncogenic genotype **(c)**, and the relative abundance of the dominant bacterial phyla **(d)**. Sample positions correspond across panels (n = 169). **(e)** Spatially resolved 16S rRNA gene sequencing of segmented small intestinal tissue pieces (5 × 5 mm) of a *Kras*^G12D^ mouse mapped onto macroscopic intestinal images. Relative abundance of Pseudomonadota, Bacillota, *Escherichia-Shigella* and *Lachnospiraceae* are shown across spatial segments encompassing tumor and adjacent non-tumorous tissue, revealing distinct microbial distribution patterns within the tumor microenvironment. **(f)** Principal coordinates analysis of shotgun metagenomic profiles derived from cecal contents of genetically engineered intestinal cancer models and co-housed WT littermates, demonstrating separation of WT, classical (*Apc*^fl/wt^) and serrated (*Braf*^V637E^, *Kras*^G12D^ and *Pik3ca*^H1047R^) CRC subtypes (*n* = 3-5; PERMANOVA, p = 0.009, R^2^ = 0.183). **(g)** Relative abundance of the dominant bacterial families identified by shotgun metagenomic sequencing (*n* = 3-5). **(h)** Principal component analysis of untargeted metabolomic profiling by reverse-phase chromatography (RPC) coupled with negative-ion liquid chromatography-mass spectrometry (LC-MS) (*n* = 3-5). **(i-k)** Quantification of primary and secondary bile acids **(i)**, short-chain fatty acids (SCFAs) **(j)**, and additional microbially derived metabolites **(k)** (*n* = 3-5; i, two-way ANOVA with Bonferroni’s multiple comparisons test, ** p < 0.01; j-k, two-way ANOVA with Dunnett’s multiple comparisons test, * p < 0.05, ** p < 0.01, and *** p < 0.001).

Due to increasing evidence implicating the gut microbiota in CRC pathogenesis,^36^ we investigated whether distinct oncogenic drivers shape intestinal microbial ecosystems during tumor development. To this end, we performed 16S rRNA amplicon sequencing of mucosa-associated microbial communities collected from different intestinal regions of genetically engineered mouse models representing major molecular routes of CRC and their wild-type (WT) littermates. Principal Coordinates Analysis (PCoA) revealed significant genotype-dependent differences in microbial composition within both the duodenum and colon prior to overt tumor formation at 8 weeks of age, indicating that epithelial oncogenic signaling influences microbial community structure before malignant transformation **(Suppl. Fig. 2a-d)**. Specifically, we observed a reduction in *Lachnospiraceae* abundance in the duodenum of both *Braf*^V637E^ and *Kras*^G12D^ mice. In addition, *Braf*^V637E^ mice exhibited increased abundances of *Bifidobacteriaceae*, *Clostridiaceae*, and *Lactobacillaceae*, whereas *Kras*^G12D^ mice showed enrichment of *Akkermansiaceae* and *Bifidobacteriaceae* **(Suppl. Fig. 2c)**. To determine how these microbial ecosystems evolve during tumor progression, we profiled mucosa-associated microbiota from tumor-bearing mice at endpoint **(Suppl. Fig. 2e-h)**. Importantly, all genotypes were randomly co-housed throughout the study to minimize cage and housing effects, which are known to strongly influence microbial composition through coprophagic transfer between animals. Despite substantial inter-individual variability, PCoA revealed a progression-associated restructuring of microbial communities. Because intestinal microbial composition differed markedly between the small and large intestine, subsequent analyses focused on the small intestine, the predominant site of tumor formation in these models **(Suppl. Fig. 1e)**. Of note, we observed a gradual, progression-dependent shift in microbial community structure from normal tissue to advanced tumors (**Fig. 1b-d**; depicted on the y-axis from bottom to top). Notably, samples from tumors and from histologically normal tissue of *Apc*^fl/wt^ mice, characterized by high tumor burden, clustered together and showed a marked enrichment of Pseudomonadota **(Fig. 1b-d)**. In contrast, WT and *Kras*^G12D^ normal tissues clustered separately and were enriched for Bacillota, while *Braf*^V637E^ tissues and tumors formed a distinct cluster characterized by increased Actinomycetota and Bacteroidota. Variance analysis across 169 small-intestinal samples confirmed that genotype and tumor stage were the dominant contributors to microbial composition, whereas sex had no detectable effect **(Suppl. Fig. 2i)**. These data demonstrate that CRC progression is accompanied by reproducible, genotype-specific microbiota signatures. To validate these findings at higher spatial resolution, we performed segmented 16S rRNA sequencing and used spatial maps for visualization. Small intestinal segments were subdivided into 5 mm × 5 mm regions, sequenced individually, and mapped onto macroscopic tissue images **(Fig. 1e, Suppl. Fig. 3a-b)**. This approach revealed a pronounced enrichment of Pseudomonadota in the immediate tumor vicinity, accompanied by increased abundance of *Escherichia*-*Shigella* and depletion of *Lachnospiraceae*. Thus, tumor-associated dysbiosis is spatially confined and directly linked to the tumor microenvironment.

We next extended our analysis to shotgun metagenomic sequencing of cecal contents from all CRC subtypes and their WT littermates. Metagenomic profiling resolved three distinct groups corresponding to WT controls, classical tumors arising in *Apc*^fl/wt^ mice, and serrated tumors driven by *Pik3ca*^H1047R^, *Braf*^V637E^, and *Kras*^G12D^ **(Fig. 1f)**. Across all tumor models, we observed a depletion of *Lachnospiraceae* and enrichment of *Enterobacteriaceae* **(Fig. 1g, Suppl. Fig. 4a)**. Beyond taxonomic differences, metagenomic analyses suggested substantial remodeling of microbial functional potential. To determine whether these changes translated into altered metabolic output, we subjected the same cecal samples to targeted and untargeted metabolomic profiling. Strikingly, metabolomic landscapes recapitulated the microbiota-based segregation of WT, classical, and serrated CRC groups **(Fig. 1h, Suppl. Fig. 4b)**. WT metabolomes remained highly similar across genotypes despite co-housing with tumor-bearing animals, whereas *Apc*^fl/wt^ tumors displayed a distinct metabolic profile and serrated subtypes clustered closely together. Consistent with these findings, bile acid metabolism was profoundly remodeled in a genotype-dependent manner, with oncogenic driver mutations reducing total bile acid abundance and altering the balance between primary and secondary bile acids **(Fig. 1i, Suppl. Fig. 4c-e)**. Beyond global shifts in bile acid abundance, analysis of cecal metabolites revealed subtype-specific remodeling of individual bile acid species and short-chain fatty acid (SCFA) profiles across CRC models. Serrated tumors were associated with increased cholic acid-derived metabolites and lactate and reduced butyrate, whereas classical tumors were characterized by marked depletion of secondary bile acids, acetate, and butyrate, revealing profound oncogene-specific rewiring of host-microbial metabolism **(Fig. 1j-k, Suppl. Fig. 4f)**.

Collectively, these results identify tumor genotype as a key organizer of microbiota structure and function, generating CRC subtype-specific host-microbial ecosystems with distinct metabolic consequences.

### A germ-free orthotopic transplantation platform enables functional interrogation of tumor-microbiota interactions

Having identified distinct genotype-associated microbial ecosystem states across endogenous CRC models, we next sought to functionally interrogate how the microbiota contributes to tumor progression. To this end, we established a modular orthotopic transplantation platform that enables controlled manipulation of microbial exposure in fully immunocompetent germ-free (GF) mice while preserving tumor formation within the native colonic microenvironment **(Fig. 2a)**. Although several colonoscopy-guided orthotopic CRC models have previously been described,^37–43^ their application under germ-free conditions has remained challenging due to the technical complexity of maintaining sterility throughout the transplantation procedure and longitudinal follow-up. To establish the platform, we generated an adenocarcinoma-derived organoid line from *Kras*^G12D^-driven tumors **(Fig. 2b)**. This model had previously been extensively characterized under SPF conditions and was therefore well suited for the development and validation of the germ-free transplantation system.^38,44,45^ Prior to transplantation, organoids were mechanically dissociated into small multicellular fragments to facilitate efficient colonoscopy-guided submucosal injection and engraftment **(Fig. 2c)**.

**Figure 2.**
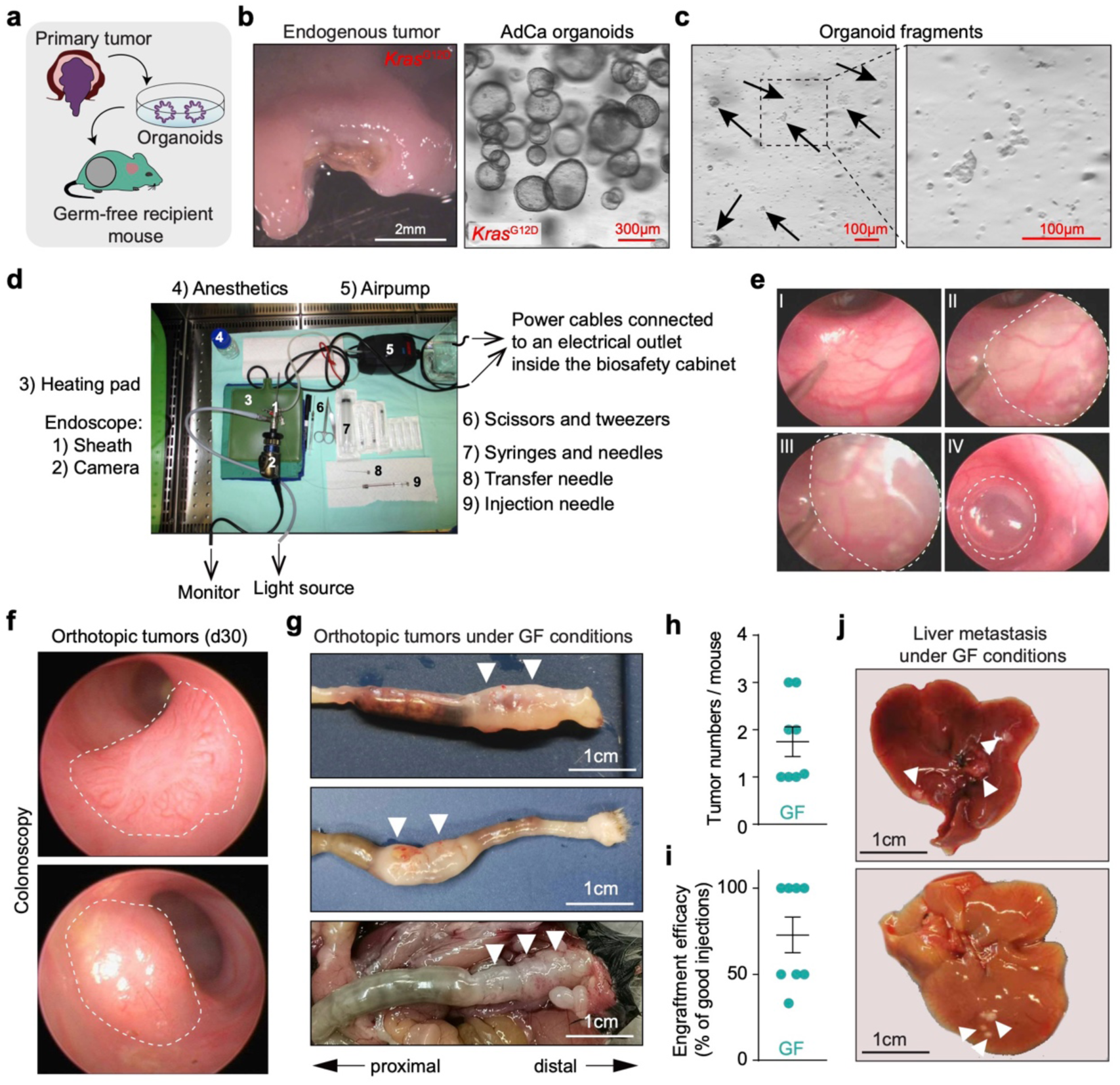
Development of a germ-free-compatible orthotopic colorectal cancer transplantation platform. **(a)** Schematic overview of the experimental workflow illustrating the generation of tumor-derived organoids from mouse models representing distinct molecular subtypes of CRC, followed by orthotopic transplantation into germ-free (GF) recipient mice. **(b)** Representative macroscopic image and hematoxylin and eosin (H&E) staining of an endogenous *Kras*^G12D^-driven adenocarcinoma used for organoid generation. **(c)** Representative brightfield images of adenocarcinoma (AdCa) organoids after mechanical dissociation into multicellular fragments for transplantation. **(d)** Schematic of the sterilized endoscopic transplantation setup adapted for use inside a germ-free isolator. All equipment required for the procedure is indicated. **(e)** Representative colonoscopic images showing submucosal injection of tumor organoids into the distal colon of a GF recipient mouse. Successful submucosal delivery results in the formation of a transient mucosal “bubble” that temporarily occludes the colonic lumen. **(f)** Representative follow-up colonoscopic image showing an orthotopic organoid-derived tumor at the injection site 30 days after transplantation. **(g)** Representative macroscopic images of the distal colon of GF recipient mice bearing orthotopic tumors following organoid transplantation. Tumors are indicated by arrowheads. **(h)** Quantification of orthotopically derived tumor burden following organoid transplantation, shown as the number of tumors per recipient mouse (*n* = 8). **(i)** Engraftment efficiency of orthotopic organoid transplantation under GF conditions, calculated as the percentage of successful tumor engraftments per technically successful submucosal injection (*n* = 8). **(j)** Representative macroscopic images of liver metastases arising from orthotopically transplanted tumors under GF conditions.

We subsequently adapted and optimized the transplantation procedure for long-term experimentation in GF animals. Briefly, all components of the endoscopic system, including the protective sheath, Hopkins optics, camera head interface, fiber-optic light cable, insufflation system and injection needles, were individually double-wrapped and sterilized by either autoclaving or vaporized hydrogen peroxide treatment before transfer into the sterilized workbench **(Fig. 2d)**. Orthotopic transplantation was subsequently performed entirely under sterile biosafety conditions using colonoscopy-guided submucosal injection of tumor organoids into the distal colon **(Fig. 2e)**. Importantly, the procedure enabled not only tumor organoid transplantation but also repeated endoscopic monitoring of tumor growth without compromising the germ-free status of the animals **(Fig. 2f)**.

To verify the robustness of the system, extensive microbiological surveillance was performed throughout all experiments using complementary culture-based and molecular approaches. Experimental animals were monitored longitudinally using aerobic and anaerobic enrichment cultures followed by plating on multiple media, including Brain Heart Infusion (BHI), Wilkins-Chalgren Anaerobe (WCA), thioglycolate, and Sabouraud media. Aerobic and anaerobic cultures were each incubated for 7 days in liquid media followed by an additional 7 days on solid media to maximize the sensitivity for detecting low-level bacterial and fungal contamination. Selected samples were further analyzed by Gram staining and broad-range bacterial 16S rDNA PCR. While sporadic contaminations occurred during the initial establishment and optimization of the procedure, iterative refinement of the workflow resulted in a robust experimental platform in which all animals included in the final analyses maintained germ-free status throughout the experimental period. These findings demonstrate that complex endoscopic orthotopic transplantation procedures can be performed reproducibly under strictly germ-free conditions.

Necropsy and histopathological analyses demonstrated efficient tumor formation with an overall engraftment efficiency exceeding 70% **(Fig. 2g-i)**, and metastatic dissemination to the liver was observed in a subset of animals in the absence of microbiota **(Fig. 2j)**.

Using this GF-compatible orthotopic transplantation platform, we next established a panel of *Kras*^G12D^-driven organoid models spanning low-grade adenoma, high-grade adenoma and adenocarcinoma states to investigate how microbial exposure influences tumor evolution **(Suppl. Fig. 5a)**. This transplantation-based approach circumvents the need for germ-free rederivation and extensive intercrossing of genetically engineered mouse models while preserving tumor growth within the native colonic microenvironment, thereby enabling systematic interrogation of microbiota-dependent effects across distinct stages of tumor progression. Integrated genomic and transcriptomic profiling revealed progressive molecular diversification along the adenoma-to-carcinoma continuum **(Suppl. Fig. 5a-b)**. Low-grade adenomas organoids (LgAd1 and LgAd2) retained largely genomically stable profiles with only focal copy-number alterations, whereas adenocarcinoma (AdCa) organoids exhibited extensive chromosomal instability characterized by broad amplifications, deletions, and complete loss of the tumor suppressor *Cdkn2a* **(Suppl. Fig. 5a-b)**. High-grade adenoma (HgAd) organoids displayed an intermediate degree of chromosomal complexity, consistent with stepwise malignant progression. Targeted sequencing further uncovered substantial inter-organoid heterogeneity. While HgAd and AdCa organoids harbored activating *Ctnnb1* mutations, LgAd-1 acquired alterations in *Trp53* and *Apc*, whereas LgAd-2 remained devoid of detectable SNVs or CNVs **(Suppl. Fig. 5a-b)**. Notably, recurrent copy-number and loss-of-heterozygosity events affected genes linked to multiple hallmark oncogenic pathways, concordant with pathway-level transcriptional changes identified by GSVA, further supporting progressive molecular evolution during tumor progression. **(Suppl. Fig. 5b)**.

Phenotypic and transcriptomic profiling revealed progressive molecular reprogramming along the adenoma-to-carcinoma continuum **(Suppl. Fig. 5b-c)**. While LgAd-2 and HgAd organoids were enriched for proliferative programs, including MYC targets, E2F signaling, and G2M checkpoint activation, AdCa and LgAd-1 organoids exhibited a shift toward inflammatory and progression-associated states characterized by TNFα/NFκB, IL6-JAK-STAT3, TGFβ signaling, apoptosis, and hypoxia. Notably, LgAd-1 organoids additionally displayed enrichment of reactive oxygen species (ROS), oxidative phosphorylation, and metabolic programs, whereas AdCa organoids showed prominent activation of epithelial-mesenchymal transition (EMT) and WNT/β-catenin signaling **(Suppl. Fig. 5b)**.

Interestingly, pERK abundance did not correlate with proliferative activity across organoid states **(Suppl. Fig. 5c)**. While LgAd-1 and LgAd-2 organoids showed the strongest MAPK pathway activation, proliferative capacity varied considerably and was highest in AdCa organoids. Together with the GSVA results, these findings suggest that progression is accompanied by diversification of the signaling programs that support tumor growth, rather than a uniform dependence on canonical ERK signaling.

### Orthotopic germ-free colorectal cancer modelling reveals stage-specific microbiota dependencies during tumor progression

Having established a panel of genetically, transcriptionally, and phenotypically distinct organoid models spanning the adenoma-to-carcinoma continuum, we next sought to determine how microbiota dependence evolves during colorectal tumor progression. To this end, we orthotopically transplanted organoids representing low-grade adenoma (LgAd), high-grade adenoma (HgAd), and adenocarcinoma (AdCa) stages into GF and SPF recipient mice, thereby enabling direct assessment of microbial contributions to tumor growth and metastatic dissemination **(Fig. 3a-d)**. As an initial benchmark, AdCa organoids engrafted efficiently under both SPF and GF conditions and exhibited comparable overall survival, demonstrating that advanced tumor states retain the capacity to grow in the complete absence of microbial signals **(Fig. 2 and 3a)**. Although liver metastases occurred more frequently in SPF recipients, metastatic dissemination was also observed in GF animals, indicating that microbiota-independent metastatic progression remains possible once malignant competence has been acquired.

**Figure 3.**
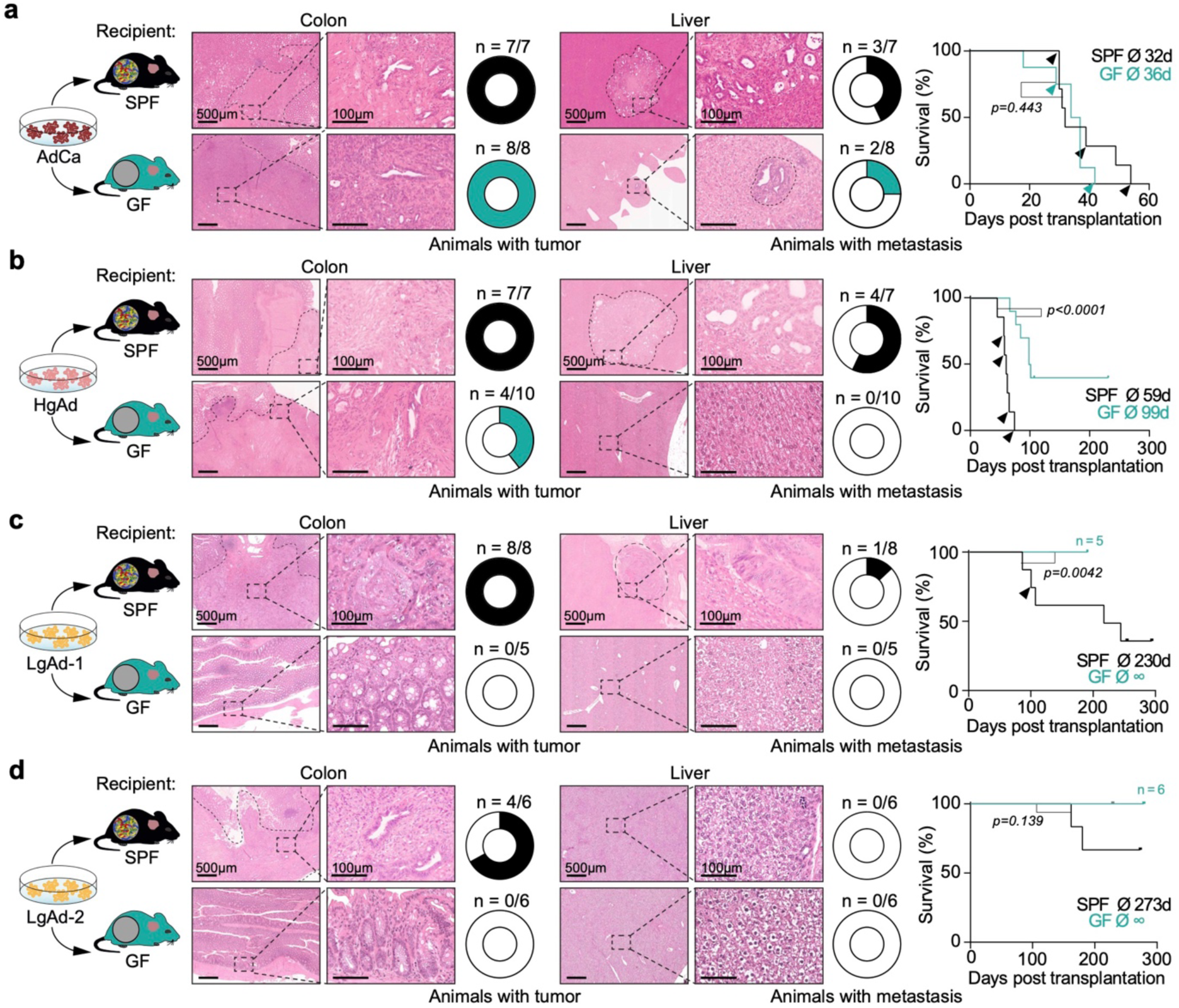
Orthotopic germ-free CRC modelling reveals stage-specific microbiota dependencies during tumor progression. **(a-d)** Representative hematoxylin and eosin (H&E)-stained sections of orthotopic primary tumors and corresponding liver metastases together with Kaplan-Meier survival analyses of recipient mice following orthotopic transplantation of adenocarcinoma (AdCa) **(a)**, high-grade adenoma (HgAd) **(b)**, low-grade adenoma 1 (LgAd-1) **(c)** and low-grade adenoma 2 (LgAd-2) **(d)** organoids into specific pathogen-free (SPF) and germ-free (GF) recipient mice. Numbers indicate mice with primary tumors or liver metastases relative to the total number of transplanted animals. Survival curves were compared using the log-rank (Mantel-Cox) test.

In striking contrast, adenoma-derived organoids displayed a profound dependence on microbial exposure. HgAd organoids reproducibly formed tumors under SPF conditions, whereas both tumor growth and metastatic dissemination were markedly impaired in GF recipients **(Fig. 3b)**. This dependency became even more pronounced in low-grade adenoma organoids. While both LgAd-1 and LgAd-2 established slowly growing tumors in SPF mice, neither line generated detectable tumors in GF recipients despite observation periods exceeding 200 days **(Fig. 3c-d)**. Longitudinal colonoscopic monitoring and endpoint histopathological analyses consistently confirmed the complete absence of engrafted primary tumors and metastatic lesions in germ-free animals.

Together, these findings reveal that microbiota independence is not a static feature of CRC but evolves during malignant progression. Early adenoma states exhibit an almost absolute requirement for microbial exposure to initiate and sustain tumor growth, whereas advanced carcinomas progressively acquire the capacity for microbiota-independent growth and dissemination. These results identify loss of microbiota dependence as a previously unrecognized hallmark of CRC progression and establish the orthotopic GF transplantation platform as a powerful framework for mechanistically dissecting host-microbial contributions to tumor evolution.

### Microbiota-dependent adenoma progression occurs in the absence of major immune or epithelial rewiring

Germ-free mice exhibit profound alterations in immune development and mucosal immune education, raising the possibility that the impaired tumor formation observed in GF recipients reflects secondary consequences of the germ-free state rather than a direct requirement for microbial signals during tumor progression.^46–48^ To address this possibility, we sought to determine whether the absence of microbiota resulted in major alterations of the immune or epithelial compartments that could account for the observed transplantation phenotypes. Because adenoma-derived organoids rarely formed tumors under GF conditions, downstream analyses were performed using *Kras^G12D^*AdCa organoids, which reproducibly engrafted in both SPF and GF recipients. Histopathological examination revealed highly similar tumor architecture across conditions **(Fig. 3)**. Multiparametric flow cytometry of tumors and adjacent colonic lamina propria demonstrated broadly comparable immune cell compositions, including total CD45^+^ leukocytes, CD3^+^, CD4^+^, CD8^+^, and CD25^+^ T cells, as well as macrophages **(Fig. 4a-b)**. Consistent with these findings, orthotopic adenocarcinomas exhibited an overall immune-desert phenotype irrespective of microbial status, and immunohistochemical quantification of CD8^+^ and FoxP3^+^ cells revealed only minimal immune infiltration in both SPF and GF tumors **(Fig. 4c-d)**. Thus, established carcinomas developed remarkably similar local immune landscapes in the presence or absence of microbiota.

**Figure 4.**
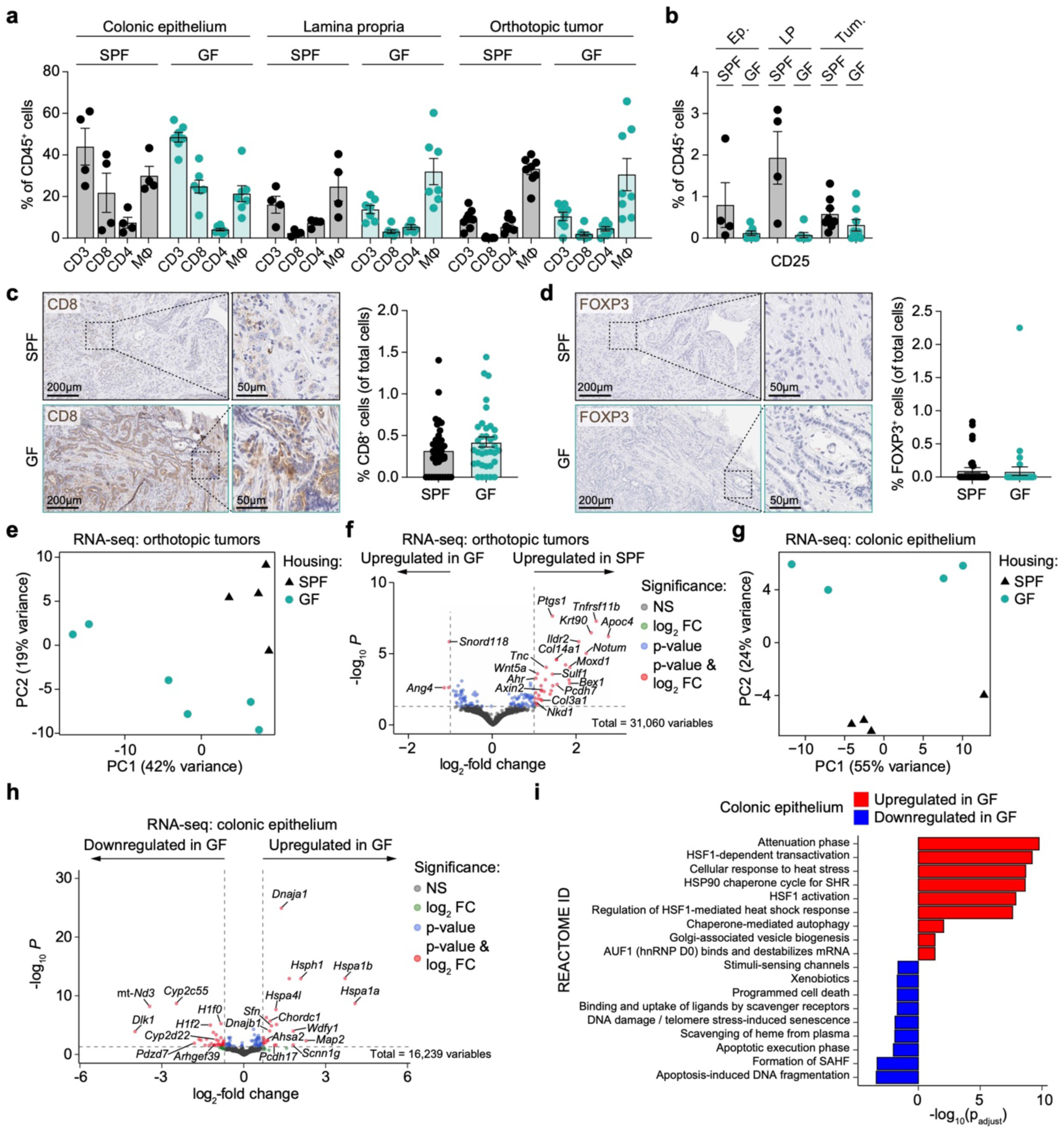
Microbiota-dependent adenoma progression occurs in the absence of major immune or epithelial rewiring. **(a-b)** Multiparametric flow cytometric analysis of immune cell populations in the colonic epithelium, lamina propria and orthotopic tumors from specific pathogen-free (SPF) and germ-free (GF) recipient mice showing the frequency of CD3^+^, CD4^+^, CD8^+^ T cells and macrophages (MΦ) among CD45^+^ cells **(a)**, and the frequency of CD25^+^ cells among CD45^+^ cells **(b)**. **(c-d)** Representative immunohistochemical staining and quantification of CD8^+^ **(c)** and FOXP3^+^ **(d)** cells in orthotopic tumors from SPF and GF recipient mice. **(e-f)** Principal component analysis (PCA) of bulk RNA sequencing data from orthotopic tumors derived from specific pathogen-free (SPF) and germ-free (GF) recipient mice **(e)** and volcano plot depicting differential gene expression analysis between tumors arising under SPF and GF conditions **(f)**. **(g-i)** Principal component analysis (PCA) of bulk RNA sequencing data from normal colonic epithelium of SPF and GF mice **(g)**, volcano plot depicting differential gene expression analysis between SPF and GF conditions **(h)**, and Reactome pathway enrichment analysis **(i)**.

To further assess whether microbial status altered tumor cell identity, we performed bulk RNA sequencing of orthotopically transplanted adenocarcinomas recovered from SPF and GF recipients. Although principal component analysis separated tumors according to microbial status **(Fig. 4e)**, differential expression analysis identified only a limited number of significantly altered genes, and no dominant transcriptional program emerged that could readily explain the profound differences in microbiota dependence observed across tumor stages **(Fig. 4f)**. We next asked whether germ-free conditions induced broader alterations in the non-transformed colonic epithelium. Bulk RNA sequencing of normal colonic epithelial tissue from age-matched SPF and GF mice likewise revealed separation by principal component analysis **(Fig. 4g)**, yet only a small number of genes were differentially expressed, with no significant enrichment of Hallmark pathways and only changes in selected Reactome pathways **(Fig. 4h-i)**. Together, these analyses provide little evidence for widespread epithelial reprogramming in the absence of microbiota.

Collectively, these findings argue against a model in which impaired adenoma growth in germ-free mice is simply a consequence of gross immune abnormalities or broad epithelial dysfunction. Instead, they support the notion that microbial signals themselves constitute a critical component of the early tumor-promoting ecosystem, whereas advanced carcinomas progressively acquire the capacity to sustain tumor growth despite the absence of microbiota.

## Discussion

CRC is increasingly recognized as a disease shaped by complex interactions between tumor-intrinsic genetic programs and the surrounding microbial ecosystem.^11,16,17^ Although numerous studies have associated specific bacterial taxa, microbial metabolites and dysbiotic community structures with colorectal tumorigenesis,^13,22,25,49^ whether distinct oncogenic drivers actively shape microbial ecosystems and how these ecosystems functionally contribute to tumor progression have remained unresolved. Here, by combining genetically engineered mouse models, longitudinal microbiota profiling and a germ-free-compatible orthotopic transplantation platform, we demonstrate that colorectal cancer genotypes are associated with distinct microbial ecosystem states and reveal a stage-dependent requirement for microbial signals during tumor progression.

A central finding of this study is that epithelial oncogenic genotype is associated with alterations in microbial community composition prior to overt tumor formation and throughout disease progression. Across multiple genetically engineered mouse models representing major molecular routes of CRC, microbial community structure segregated according to tumor genotype and progression stage despite random co-housing of animals of the different mouse lines. These ecosystem differences were not limited to taxonomic composition but extended to metagenomic functional capacity and fecal metabolomic profiles, suggesting that distinct oncogenic programs establish specific host-associated ecological niches. Notably, across all tumor models, we consistently observed depletion of members of the *Lachnospiraceae* family and enrichment of *Enterobacteriaceae*, indicating the existence of conserved ecological responses to neoplastic transformation that transcend specific oncogenic drivers. The depletion of the *Lachnospiraceae* family is particularly intriguing, as recent large-scale meta-analyses of human CRC microbiome datasets have identified *Lachnospira*, a prominent member of the *Lachnospiraceae* family, as one of the taxa most consistently and strongly negatively associated with CRC across diverse patient cohorts, age groups, and sequencing methodologies, suggesting that our findings capture fundamental features of host-microbiota interactions during colorectal tumorigenesis.^50^ In addition, these findings are consistent with emerging evidence that epithelial signaling pathways can influence microbial community assembly through alterations in barrier function, nutrient availability, immune signaling and epithelial metabolism.^17,51,52^ Together, these observations support a model in which oncogenic programs not only shape tumor cell-intrinsic properties but also actively remodel the surrounding microbial ecosystem, establishing a bidirectional relationship whereby tumor-induced ecological changes may subsequently influence tumor behavior. Importantly, these ecosystem-level alterations extended beyond taxonomic composition to microbial functional capacity and metabolite production. Distinct CRC genotypes exhibited characteristic metagenomic and metabolomic profiles, including remodeling of bile acid and short-chain fatty acid metabolism. The stage-specific microbiota dependencies revealed in our transplantation experiments likely emerged from broader ecological and metabolic interactions between tumor cells and their resident microbiota.

A major obstacle to mechanistic interrogation of tumor-microbiota interactions has been the lack of experimental systems that combine defined tumor genotypes with precise microbial manipulation under physiologically relevant conditions. Germ-free studies using genetically engineered mouse models are technically demanding, require extensive breeding and rederivation efforts, and are frequently confounded by variability in tumor onset and progression. By adapting colonoscopy-guided orthotopic transplantation for long-term germ-free experimentation, we establish a platform that overcomes many of these limitations. Importantly, this approach enables direct side-by-side comparison of the same genetically and molecularly defined tumor line in the presence or absence of microbiota while preserving growth within the native colonic microenvironment. Such experimental control is difficult to achieve in endogenous mouse models and impossible in human cohorts, where microbial exposure, host genetics and tumor evolution are inherently intertwined. The ability to compare identical tumor genotypes across distinct microbial environments therefore provides a powerful framework for establishing causal relationships between microbial exposure and tumor progression.

While numerous studies have implicated the microbiota in colorectal tumorigenesis, most have examined its contribution to tumor initiation or overall tumor burden rather than its role across distinct stages of disease progression.^11,13,14^ Our findings instead suggest that microbiota dependence itself evolves during malignant progression. Across the progression continuum analyzed here, microbiota dependence decreased as tumors acquired increasingly advanced molecular features. Low-grade adenomas exhibited an almost absolute requirement for microbial exposure, high-grade adenomas retained partial dependence, whereas adenocarcinomas remained capable of sustained growth and metastatic dissemination in the complete absence of microbiota. Although additional models will be required to determine the generality of this phenomenon, these observations raise the possibility that progressive acquisition of microbiota independence represents a previously underappreciated feature of advanced CRC evolution.

This concept is compatible with current models of malignant progression. Early neoplastic lesions remain strongly dependent on environmental signals that promote epithelial proliferation, survival, and metabolic adaptation.^53,54^ A growing body of evidence indicates that microbial metabolites, inflammatory mediators, and microbe-host signaling pathways can provide such cues during early tumorigenesis.^14–16,49^ In contrast, advanced tumors accumulate genetic, epigenetic, and transcriptional alterations that may progressively substitute for exogenous growth-promoting signals.^53–55^ Notably, the loss of microbiota dependence observed here coincided with increasing genomic complexity and the emergence of inflammatory, hypoxic, and mesenchymal-like transcriptional programs. While the specific molecular determinants underlying this transition remain to be defined, our data indicate that microbiota dependence decreases in parallel with increasing tumor complexity and the acquisition of advanced transcriptional states during malignant progression. Together, these findings suggest that malignant progression is accompanied by a shift from ecosystem dependence toward tumor cell autonomy. More broadly, our results imply that the impact of the microbiota on tumor progression is likely to depend critically on the intrinsic degree of tumor autonomy, raising the possibility that experimental models representing distinct stages of tumor evolution may yield fundamentally different conclusions regarding microbiota dependence. Defining these stage- and genotype-specific dependencies more comprehensively will therefore be essential for understanding when, and in which tumors, microbial signals constitute actionable determinants of cancer progression.

An important alternative explanation for our findings relates to the well-established immunological abnormalities of GF animals. GF mice exhibit altered lymphoid organ development, impaired immune education and defects in both innate and adaptive immune maturation.^46–48,56^ It could therefore be argued that differences in tumor progression simply reflect differences in immune competence rather than direct microbiota-dependent effects. However, several observations suggest that broad immune or epithelial alterations are unlikely to fully account for the observed phenotype. Comprehensive immunophenotyping revealed highly similar immune landscapes in SPF and GF adenocarcinomas, which were characterized by an overall immune-desert phenotype irrespective of microbial status. Likewise, bulk transcriptomic profiling identified only limited differences in tumor and epithelial gene expression despite clear separation by principal component analysis. Collectively, these findings provide little evidence for major immune rewiring or baseline epithelial dysfunction that could readily explain the profound impairment of adenoma growth observed under germ-free conditions.

Indeed, one might predict the opposite outcome if impaired immune education were the dominant determinant of tumor growth. Because GF mice possess less mature tumor-surveillance mechanisms, reduced adaptive immune priming and impaired inflammatory responses, diminished immune-mediated tumor control would be expected to facilitate rather than impair tumor establishment. Instead, adenoma-derived organoids failed to form tumors in GF recipients despite the presence of a comparatively less educated immune system. This observation supports the notion that microbially derived signals themselves provide critical functions required for early tumor establishment and progression. Although the precise nature of these signals remains unresolved, they may include microbial metabolites, inflammatory mediators, microbial-associated molecular patterns or indirect effects on stromal and epithelial signaling networks. Importantly, the stage-specific microbiota dependence uncovered by our germ-free-compatible orthotopic transplantation platform provides a clear rationale and an experimentally tractable framework to dissect the molecular and cellular mechanisms through which microbial signals promote early colorectal tumor progression.

Recent work has increasingly implicated the microbiota as a determinant of therapeutic response and disease outcome across multiple cancer types.^18,19,57,58^ Our study extends these observations by suggesting that the importance of microbial signals may depend strongly on tumor stage. If confirmed in additional models and human disease, these findings could have important implications for microbiota-targeted interventions. Early lesions may be particularly susceptible to strategies that disrupt tumor-promoting microbial functions, whereas advanced carcinomas may have already acquired mechanisms conferring progressive independence from microbial support. The orthotopic germ-free platform described here provides a versatile experimental framework to address these questions through controlled colonization experiments, defined microbial consortia, fecal microbiota transplantation and systematic interrogation of microbial metabolites.

In summary, we identify epithelial oncogenic genotype as a major determinant of CRC-associated microbial ecosystem structure and function and establish a germ-free-compatible orthotopic transplantation platform for causal investigation of tumor-microbiota interactions. Using this system, we demonstrate that microbiota dependence evolves during colorectal tumor progression, with early adenomas exhibiting an almost absolute requirement for microbial exposure and advanced carcinomas progressively acquiring microbiota-independent growth properties. Together, these findings support a model in which colorectal tumor evolution is accompanied by a transition from ecosystem dependence toward tumor cell autonomy and establish microbiota independence as a previously underappreciated feature of advanced disease.

## Methods

### Animal experiments

All animal experiments were approved by the relevant governmental authorities (Regierung von Oberbayern) and conducted in accordance with institutional, national and European regulations for animal welfare. Mice were maintained on a C57BL/6 background under specific pathogen-free (SPF) conditions at the ZPF animal facility (TUM University Hospital, Technical University of Munich, Munich, Germany). Germ-free C57BL/6 mice used for transplantation experiments were bred, housed and experimentally manipulated in isolators or gnotobiotic cages (IsoCages P, Tecniplast) at the Core Facility Gnotobiology, ZIEL Institute for Food & Health (Technical University of Munich, Freising, Germany).

To generate intestinal cancer models, mice carrying the intestinal epithelial-specific Villin-Cre transgene ^59^ were crossed with mice harboring *Kras*^G12D^, *Braf*^V637E^, *Pik3ca*^H1047R^ or *Apc* alleles ^31,32,34,35^ as indicated for the individual experiments. Both female and male mice were included. To minimize confounding effects arising from genotype-specific breeding line microbiota, mice of different genotypes were randomly distributed across cages after weaning and subsequently co-housed whenever possible. This strategy promotes microbiota normalization through coprophagic behavior and reduces cage- and lineage-associated microbiota effects. For transplantation experiments, recipient mice were randomly assigned to experimental groups. Animals were monitored throughout the study and euthanized upon reaching predefined humane endpoints. At necropsy, the intestinal tract and major metastatic sites, including liver, lung and regional lymph nodes, were examined macroscopically and histologically.

### Microbiological monitoring of germ-free mice

Germ-free mice were derived from germ-free breeding colonies and monitored microbiologically before and during experiments. Breeding isolators were tested for sterility after each opening by culture, Gram staining, and qPCR. In addition, filter samples were collected every six months for PCR-based screening (PRIA) for microorganisms included in the FELASA recommendations. For experimental animals, microbiological monitoring was performed using culture-based methods. Fecal pellets or swab samples were collected and tested for aerobic and anaerobic microbial growth. For aerobic culture, samples were inoculated into 50 mL Brain Heart Infusion (BHI) broth in 250 mL Schott bottles and incubated at 37°C for 7 days with shaking at 180 rpm. Subsequently, 50 μL of the enrichment culture was plated onto BHI agar and incubated aerobically at 37°C for an additional 7 days. For anaerobic culture, all procedures were performed in an anaerobic chamber (4.5% H_2_, 10% CO_2_, 85.5% N_2_). Samples were inoculated into 9 mL degassed Wilkins-Chalgren Anaerobe (WCA) broth in Hungate tubes and incubated at 37°C for 7 days with shaking at 180 rpm. Subsequently, 50 μL of the enrichment culture was plated onto WCA agar and incubated anaerobically at 37°C for an additional 7 days. Routine microbiological testing also included incubation in thioglycolate broth and on Sabouraud agar for one week, followed by plating of thioglycolate cultures onto BHI and WCA agar and incubation for an additional week. Gram staining and broad-range bacterial PCR analyses were performed on selected samples. Microbiological analyses were performed at two locations, either internally at the Core Facility Gnotobiology or independently by the Core Facility Microbiome. Samples with microbial growth were re-evaluated using repeat culture and/or complementary methods, including Gram staining and PCR. Germ-free status was considered compromised only when microbial contamination was confirmed by independent testing or by concordant results obtained using multiple microbiological assays.

For taxonomic identification of cultured microorganisms, colonies were suspended in molecular biology-grade water and subjected to 16S rRNA gene amplification using primers 27F (5′-AGAGTTTGATCCTGGCTCAG-3′) and 1492R (5′-GGTTACCTTGTTACGACTT-3′). PCR reactions contained 25 μL 2× DreamTaq PCR Master Mix (Thermo Scientific), 2.5 μL of each primer (10 μM), 10 μL template suspension, and nuclease-free water to a final volume of 50 μL. Cycling conditions were 95°C for 6 min; 30 cycles of 95°C for 30 s, 55°C for 20 s, and 72°C for 90 s; followed by a final extension at 72°C for 10 min. PCR products were verified by agarose gel electrophoresis, purified using the QIAquick PCR Purification Kit (Qiagen), and subjected to Sanger sequencing (Eurofins Genomics).

### Tumor cell isolation

To generate tumor-derived organoids, tumor tissues were minced with a scalpel and incubated in PBS containing 10% PenStrep for 15 min on ice. The tissues were then washed and digested with 230 U of collagenase type IV (Merck) at 37°C for 30 min. The digest was filtered through a 100 μm strainer, washed with PBS, and the collagenase was inactivated by incubation with 10% FCS for 10 min. Following centrifugation at 300 × g for 5 minutes, approximately 100 cell clumps were resuspended in 50 μL of Matrigel (Corning) and plated in a 24-well plate.

### Organoid culture

The organoid cultures were maintained in a conditioned medium (50% diluted) containing Wnt3A, R-spondin, and Noggin, produced by L-WRN cells (ATCC® CRL-3276™), supplemented with 10 μM Y-27632 (STEMCELL Technologies). The medium was refreshed every 2-3 days, and cultures were passaged every 4-6 days. For passaging, organoid cultures were incubated with Cell Recovery Solution (Corning) for 30 min on ice to facilitate the dissolution of Matrigel, followed by centrifugation at 300 × g for 5 min. Cells were then enzymatically digested with TrypLE™ Express Enzyme (Thermo Fisher Scientific) diluted 1:1 in PBS and incubated at 37°C for 3 min. After washing once with PBS and centrifugation at 300 × g, cells were resuspended in Matrigel (Corning) and plated in a 24-well plate (50 μL/well). Organoid cultures were routinely tested for mycoplasma by PCR analysis. Organoid line authentication was performed by re-genotyping of engineered alleles from genomic DNA (gDNA).

### Formalin-fixation and paraffin-embedding (FFPE) of organoids

For each paraffin embedding cassette, organoids from at least three confluent wells were pooled. For FFPE preparation, organoids were cultured in 24-well plates on sterile 13-mm round glass coverslips (Carl Roth). After 4 days of culture, the medium was removed and organoids were fixed with 500 μL formalin for 45 min at room temperature. Following fixation, formalin was aspirated, and the organoids were gently detached from the coverslips and transferred into embedding cassettes (Bionet, Engelbrecht). Samples were stored in PBS until subsequent dehydration and paraffin embedding.

### Histopathological assessment

FFPE tissues and organoids were cut into 2 µm sections with a microtome. Tissue slices were subjected to deparaffinization and dehydration prior to histopathological analysis. For hematoxylin and eosin (H&E) staining, experiments were subsequently conducted following standard protocols. Histopathological assessment was performed by experienced gastrointestinal pathologists (M.J., T.G. and K.S.). Tumor grade was determined based on gland formation, with tumors categorized as low-grade (≥50% gland formation) or high-grade (<50% gland formation).^60^ The presence of metastatic disease was additionally assessed in the liver, lung, pancreas and spleen.

Immunohistochemistry (IHC) was performed on formalin-fixed paraffin-embedded (FFPE) tissue sections at the Comparative Experimental Pathology (CEP) facility of TUM University Hospital using a Bond RXm automated staining platform (Leica Biosystems). Sections were deparaffinized and subjected to heat-induced epitope retrieval using Bond Epitope Retrieval Solution 1 (ER1; citrate buffer, pH = 6.0) or Solution 2 (EDTA-based buffer, pH = 9.0) for 30 min. Endogenous peroxidase activity was blocked prior to incubation with primary antibodies against CD8 (clone 309, Sino Biological, 1:100), FOXP3 (clone FJK-16s, Thermo Fisher Scientific, 1:100), Ki67 (clone SP6, Abcam, 1:50) and pERK (clone 20G11, Cell Signaling, 1:1,000) for 15 min at room temperature. Antibody detection was performed using the Bond Polymer Refine Detection Kit (Leica Biosystems) without the post-primary reagent and visualized using 3,3′-diaminobenzidine (DAB), followed by hematoxylin counterstaining. Whole-slide images were acquired using a Leica Aperio AT2 slide scanner.

Quantification of IHC staining was performed in QuPath (v0.7.0). Three tumor regions were annotated per animal, and six regions of interest (ROIs) were analyzed within each region. Positive cell detection and tissue segmentation were performed using automated image analysis with customized DAB optical density thresholds. The proportion of positively stained cells was determined for each marker.

### Flow cytometry

Freshly harvested tumor samples were enzymatically digested using the same protocol as described for tumor organoid isolation. Following digestion, cell pellets were resuspended in PBS containing 2% FCS. Cells were incubated on ice for 10 min with anti-mouse CD16/CD32 Fc Block (BioLegend, 1:100) and subsequently stained with Zombie Aqua Fixable Viability Dye (BioLegend, 1:500). For adaptive and innate immune cell profiling, cells were stained with antibodies against CD45 (clone I3/2.3), CD3ε (clone 145-2C11), CD4 (clone GK1.5), CD8a (clone 53-6.7), CD25 (clone PC61) and F4/80 (clone BM8). Antibodies were purchased from BioLegend or BD Biosciences. Samples were acquired on a BD LSRFortessa flow cytometer, collecting up to 1 × 10^6^ events per sample. Data were analyzed using FlowJo software v10.6.2 (BD Biosciences).

### Orthotopic organoid transplantation

Orthotopic transplantations of organoids were performed in mice housed under specific pathogen-free (SPF) conditions at the ZPF animal facility (TUM University Hospital, Technical University of Munich, Munich, Germany), as previously described.^38,44,61^ Briefly, organoids were dissociated into clusters of 5-10 cells and resuspended in a transplantation medium consisting of PBS, 1× B27, 1× N2, L-glutamine (all from Gibco, Thermo Fisher Scientific), 10% Matrigel (Corning) and 10 µM Y-27632 (STEMCELL Technologies). For each injection (2-3 per mouse), 50 dissociated organoids were prepared in a volume of 100 µL. The procedure involved anesthetizing the mice, followed by gently rinsing the colon with PBS using a syringe and a straight oral gavage needle. Colonoscopy was performed using a rigid endoscope from Karl STORZ (Ø 1.9 mm) with linear Hopkins lens optics (ColoView System). Organoids were injected into the submucosa of the distal colon and rectum using a flexible fine needle (Hamilton; 33-gauge, custom length of 16 inches, custom point style of 4 at 45°). Correct submucosal injections were identified by the formation of a bubble that occluded the intestinal lumen. A scoring system was employed to correlate the quality of injections with the experimental outcomes.

All animals with signs of sickness were sacrificed in compliance with the European guidelines for the care and use of laboratory animals. For necropsy of tumor-bearing mice, the large intestine was macroscopically checked for the presence of primary tumors and metastases at the main metastatic sites (liver, lung, lymph nodes).

### Orthotopic colonic organoid transplantation under germ-free conditions

Orthotopic colonic organoid transplantation into germ-free mice was performed in the Core Facility Gnotobiology of the ZIEL Institute for Food & Health. Prior to transplantation, all surgical instruments, consumables, and endoscopic equipment were sterilized by autoclaving or vaporized hydrogen peroxide decontamination, as appropriate. Sterility was maintained throughout all procedures using established gnotobiotic handling practices.

Organoids were prepared as described above for orthotopic transplantation and transported on ice to the gnotobiotic facility. Orthotopic transplantation was performed using the same endoscopic submucosal injection procedure, instrumentation, and transplantation site (distal colon and rectum) as described for SPF-housed mice, with all manipulations conducted under aseptic germ-free conditions. Only animals that remained free of detectable microbial contamination throughout the study were included in downstream analyses (see above).

### Genomic DNA and RNA isolation of organoids

Three densely-populated wells of organoids in a 24-well culture plate were used for the isolation of either genomic DNA (gDNA) using the GenElute Mammalian Genomic DNA Miniprep Kit (Sigma-Aldrich) or RNA using the RNeasy Mini kit (Qiagen) according to the manufacturer’s instructions. Briefly, organoids were grown for up to 3-4 days in Matrigel in a 24-well plate. On the day of cell harvest, cultures were incubated with Cell Recovery Solution to separate the Matrigel from the cellular fraction. Cell pellets were snap frozen in liquid N_2_ and stored at -80°C for downstream applications. For RNA isolation, organoids were lyzed in 350 μL RLT buffer containing β-mercaptoethanol. gDNA and RNA concentrations were determined using a Qubit fluorometer (Thermo Fisher Scientific).

### RNA isolation of intestinal tissues and tumors

The intestine was dissected, cleaned, flushed with ice-cold PBS, longitudinally opened, everted, and the epithelium was scraped off using a glass slide. The epithelial scrapings and tumor pieces were collected, snap-frozen in liquid N_2_, and stored at -80°C for RNA isolation. The tissues were lysed and homogenized in tubes filled with ceramic beads (Ø 1 mm, Carl-Roth) and 600 μL RLT buffer containing β-mercaptoethanol using the Precellys Evolution (Bertin Technologies). RNA isolation was performed using the RNeasy Mini Kit (Qiagen) according to the manufacturer’s protocol.

### RNA-sequencing (RNA-seq) and analysis of organoids and tumor tissues

Library preparation for bulk-sequencing of poly(A)-RNA was performed as described previously ^62^. Briefly, barcoded cDNA of each sample was generated with a Maxima RT polymerase (Thermo Fisher Scientific) using an oligo-dT primer containing barcodes, unique molecular identifiers (UMIs) and an adaptor. Ends of the cDNAs were extended by a template switch oligo (TSO) and full-length cDNA was amplified with primers binding to the TSO-site and the adaptor. NEBNext Ultra II FS kit was used to fragment cDNA. After end repair and A-tailing, a TruSeq adapter was ligated, and 3’-end-fragments were finally amplified using primers with Illumina P5 and P7 overhangs. In comparison to Parekh et al.^62^ the P5 and P7 sites were exchanged to allow sequencing of the cDNA in read1 and barcodes and UMIs in read2 to achieve a better cluster recognition. The library was sequenced on a NextSeq 550 (Illumina) with 63 cycles for the cDNA in read1 and 16 cycles for the barcodes and UMIs in read2. Analysis of RNA-seq data was performed as described previously.^45,63^ Briefly, sequencing reads were demultiplexed, adapter and poly(A) sequences were trimmed, and reads were aligned to the mouse reference genome. Unique molecular identifiers (UMIs) were used to collapse PCR duplicates and generate gene-level count matrices for downstream analyses. The resulting UMI count tables were subsequently used for normalization, differential expression, and pathway enrichment analyses.

### Low-coverage whole genome sequencing (lcWGS) and copy number variant (CNV) analysis

Low-coverage whole genome sequencing (lcWGS) was performed using 200 ng of gDNA from organoids and matched normal samples from mouse tail biopsies when available. Libraries were prepared using the TruSeq DNA Nanokit (Illumina) following the manufacturer’s instructions. Resulting libraries were analyzed on a 2100 Bioanalyzer instrument (Agilent Technologies) and sequenced on a NextSeq 550 (Illumina) or NovaSeq 6000 (Illumina) system. Copy-number variant analysis was performed as described previously.^45,63^ Briefly, lcWGS reads were quality-filtered, aligned to the mouse reference genome, and processed to remove duplicates and generate analysis-ready BAM files. Copy number profiles were inferred using CNVKit by comparing normalized sequencing coverage across genomic regions to a reference baseline, allowing the identification and visualization of genome-wide copy number alterations.

### DNA isolation of intestinal tissues, tumors and cecal stool samples

To characterize mucosa-associated microbial communities, bulk tumors or longitudinally opened intestinal tissue segments (∼0.5 cm) were snap-frozen in liquid N_2_ immediately after collection. For analysis of luminal microbiota, cecal stool samples were collected and snap-frozen in liquid N_2_. Genomic DNA was isolated from all samples using the Maxwell® RSC Fecal Microbiome DNA Kit on a Maxwell® RSC Instrument (Promega) according to the manufacturer’s instructions.

### 16S rRNA gene sequencing and microbial profiling

16S rRNA gene sequencing of DNA isolated from bulk tumors or whole tissue pieces was performed as previously described.^64^ Briefly, the V3-V4 hypervariable regions of the bacterial 16S rRNA gene were amplified using primers 341F (5′-CCTACGGGNGGCWGCAG-3′) and 785R (5′-GACTACHVGGGTATCTAATCC-3′) in a two-step PCR workflow. Amplicons were purified using AMPure XP magnetic beads (Beckman Coulter), indexed, pooled and sequenced on an Illumina MiSeq platform using v3 chemistry (2 × 300 bp).

Raw sequencing reads were processed using the IMNGS pipeline.^65^ Reads were quality filtered, merged and dereplicated, and chimeric sequences were removed to generate zero-radius operational taxonomic units (zOTUs). Taxonomic assignment was performed against the SILVA v138.2 database using Taxonomy-Informed Clustering (TIC).^66^ Downstream analyses were conducted using the NAMCO Shiny App,^67^ built on the Rhea workflow.^68^

The zOTU count table was normalized by total-sum scaling to the sample with the lowest sequencing depth. Taxonomic profiles were aggregated to genus level using the Rhea binning module. Genera detected in at least 10% of samples and exhibiting a mean relative abundance ≥0.1% were retained for taxon-level analyses. Alpha diversity was assessed using observed richness and Shannon effective diversity (exp(H’)). Beta diversity was calculated using generalized UniFrac distances based on a midpoint-rooted maximum-likelihood phylogenetic tree and visualized by principal coordinates analysis (PCoA). Community differences were assessed by permutational multivariate analysis of variance (PERMANOVA; adonis2, 999 permutations), and homogeneity of dispersion was evaluated using PERMDISP (betadisper, vegan package).

### Spatial 16S rRNA mapping and downstream analysis

Spatial 16S rRNA maps were generated by manually segmenting tissue images using Inkscape 13.0. Images were imported into a standard SVG template and individual grid regions were traced as closed polygon paths using the Bezier tool. Each segmented region was assigned a unique grid identifier corresponding to the sampling location, and group IDs were verified in the XML editor to ensure compatibility with downstream spatial plotting. Downstream 16S analyses were performed in R using a custom script based on the spatialHeatmap package.^69^ Normalized sub-operational taxonomic unit (sOTU) abundance tables, sample metadata, taxonomy tables and alpha-diversity output were calculated as described above and imported into the R package for each sample. Grid identifiers in the sequencing tables were matched to the corresponding SVG segment IDs. Spatial heatmaps were then generated for selected sOTUs, total bacterial abundance and alpha-diversity metrics, including effective Shannon diversity and effective richness. Heatmaps were exported as PDF files, with spatial color gradients overlaid onto the segmented tissue maps to visualize the distribution of microbial taxa and diversity across the sampled tissue regions.

### Shotgun metagenomic sequencing of cecal stool samples and microbial profiling

Genomic DNA isolated from cecal stool samples was used for shotgun metagenomic sequencing. Sequencing libraries were prepared using the TruSeq® DNA Library Preparation Kit (Illumina) according to the manufacturer’s protocol. Briefly, genomic DNA was randomly fragmented, end repaired, A-tailed and ligated to Illumina sequencing adapters, followed by size selection and PCR amplification. Library quality and fragment size distribution were assessed using a 2100 Bioanalyzer (Agilent Technologies), and DNA concentrations were determined by Qubit fluorometry and quantitative PCR. Libraries were pooled and sequenced on an Illumina NovaSeq platform in paired-end (PE) 150 bp mode (2 × 150 bp), targeting approximately 12 Gb of sequencing data per sample after demultiplexing.

Raw sequencing reads were quality filtered and adapter trimmed using BBduk (v38.91). Low-quality bases were removed from both read ends (qtrim = rl, trimq = 3), reads with an average quality score below 25 were discarded, adapter sequences were removed (ktrim = r, k = 23, mink = 11, hdist = 1, tpe = true, tbo = true), and reads shorter than 50 bp were excluded. Taxonomic profiling was performed using the mOTUs4 pipeline,^70^ with sequencing reads mapped to the Genome Taxonomy Database (GTDB).

For community-level analyses, metagenomic operational taxonomic unit (mOTU) abundance profiles were rarefied to 10,000 reads per sample using the rrarefy function from the vegan R package (v2.7). Bray-Curtis dissimilarities were calculated using vegdist and visualized by principal coordinates analysis (PCoA). Differences in microbial community composition between genotypes were assessed by PERMANOVA (adonis, 999 permutations).

For differential abundance analyses, pairwise comparisons between each experimental condition and wild-type (WT) control mice were performed using the MaAsLin2 R package.^71^ Prior to statistical testing, low-abundance and low-prevalence mOTUs were filtered, retaining taxa with a relative abundance ≥10^-3^ in at least two WT mice and at least two mice of any mutant genotype. Differential abundance was assessed using linear regression models, with effect sizes reported as estimated regression coefficients (MaAsLin2 coef values). Statistical significance was determined using coefficient-specific Wald t-tests, and p-values were adjusted for multiple hypothesis testing using the Benjamini-Hochberg false discovery rate (FDR) procedure. Associations with FDR-adjusted p-values < 0.05 were considered statistically significant. For phylogenetically informed visualization, mean log_2_ fold changes in relative abundance between each genotype and WT controls were calculated for all retained mOTUs and displayed as heatmaps alongside a phylogenetic tree generated using iTOL.70.^72^

### Sample preparation for metabolomic profiling of cecal stool

Approximately 20 mg of cecal content was transferred into 2 mL bead-beating tubes (FastPrep Matrix D, MP Biomedicals). Samples were extracted in 1 mL methanol-based extraction solvent containing dehydrocholic acid (1.3 µmol/L) as an internal standard to control for extraction efficiency and sample processing losses. Mechanical lysis and metabolite extraction were performed using a FastPrep-24™ 5G instrument (MP Biomedicals) equipped with a CoolPrep™ adapter maintained on dry ice. Samples were subjected to three extraction cycles of 20 s at 6 m s^-1^ with intermittent cooling periods. Following extraction, samples were processed according to established metabolomics workflows and subjected to untargeted mass spectrometric analysis.

### Untargeted metabolomic profiling

Untargeted metabolomic analyses were performed on a Nexera UHPLC system (Shimadzu) coupled to a TripleTOF 6600 quadrupole time-of-flight mass spectrometer (AB Sciex). To maximize metabolome coverage, samples were analyzed using both hydrophilic interaction liquid chromatography (HILIC; BEH Amide, Waters) and reversed-phase chromatography (Kinetex XB-C18, Phenomenex). Chromatographic separations were performed using established solvent gradients with ammonium acetate-based mobile phases for HILIC analyses and formic acid-modified aqueous and acetonitrile mobile phases for reversed-phase analyses. Samples were injected in randomized order and analyzed in both positive and negative ionization modes using information-dependent acquisition (IDA). Quality control (QC) samples generated by pooling aliquots from all study samples were injected at regular intervals throughout the analytical sequence to monitor instrument stability and analytical reproducibility. Raw data were converted to mzXML format using ProteoWizard and processed in R using the XCMS package.^73,74^ Peaks were detected using the matchedFilter algorithm, aligned across samples, and grouped into metabolite features based on retention time and mass-to-charge ratio (*m/z*). Metabolite annotation was performed using accurate mass measurements and MS/MS fragmentation spectra through comparison with entries in the Human Metabolome Database (HMDB), the Global Natural Products Social Molecular Networking (GNPS) database, publicly available spectral libraries accessed through MS-DIAL,^75^ and BayBioMS internal reference standards. Annotation confidence was assessed using a four-point MS2 scoring system, and only metabolites with MS2 scores of 3 (spectral match to a public database) or 4 (spectral match to an internal reference standard) were retained for downstream analyses. Metabolite abundances were quantified as integrated peak areas of aligned metabolite features. Retention time correction was performed using peaks consistently detected across samples.

For downstream analyses, metabolite intensities were log_10_-transformed prior to statistical analysis. Principal component analysis (PCA) was performed using the prcomp function. Missing values were imputed using half-minimum value replacement on a per-feature basis, metabolites exhibiting zero variance were excluded, and data were mean-centered and scaled to unit variance before PCA. For heatmap visualization, metabolite-wise Z-scores were calculated by centering each metabolite intensity to its mean abundance across all samples and scaling by the corresponding standard deviation. Genotype-specific metabolic profiles were generated by averaging metabolite Z-scores across samples within each genotype group, resulting in a genotype-averaged Z-score matrix. The matrix was visualized using the ComplexHeatmap package,^76^ with metabolites hierarchically clustered according to abundance patterns and grouped according to their biochemical class.

### Targeted bile acid quantification

Targeted bile acid profiling was performed using stable isotope dilution mass spectrometry. Briefly, isotopically labeled bile acid standards were added to sample extracts prior to analysis. Bile acids were quantified using an ExionLC AD ultra-high-performance liquid chromatography (UHPLC) system coupled to a QTRAP 5500 triple quadrupole mass spectrometer (Sciex). Data acquisition was performed in multiple reaction monitoring (MRM) mode as previously described by Reiter et al.^77^ Instrument control and data acquisition were performed using Analyst v1.7 software (Sciex). Targeted data underwent analysis using MetaboAnalyst and GraphPad Prism v10.1.1.

### Targeted short-chain fatty acid quantification

Short-chain fatty acids (SCFAs) were quantified using 3-nitrophenylhydrazine (3-NPH) derivatization as previously described.^78^ Briefly, cecal extracts were spiked with isotopically labeled internal standards and derivatized using 3-NPH and EDC chemistry prior to analysis. Derivatized SCFAs were separated by reversed-phase UHPLC on a Kinetex C18 column (Phenomenex) and quantified using an ExionLC AD UHPLC system coupled to a QTRAP 5500 triple quadrupole mass spectrometer (Sciex) operating in multiple reaction monitoring (MRM) mode. Quantification was performed using calibration standards and isotope-labeled internal controls. Instrument control and data acquisition were performed using Analyst v1.7 software (Sciex). Targeted data underwent analysis using MetaboAnalyst and GraphPad Prism v10.1.1.

### Statistical analysis

GraphPad Prism v10.1.1 was used for graphical representation and statistical analysis. The data are presented as mean values ± SEM. Survival comparison was performed with a log-rank (Mantel-Cox) test. Differences in means among experimental groups were assessed through either ordinary one-way or two-way analysis of variance (ANOVA) for multiple comparisons (Tukey’s, Bonferroni’s, or Dunnett’s correction), as indicated in the respective figure legends. Fisher’s exact test was used to compare the proportions of injections resulting in engraftment versus no engraftment. Significance was predetermined at p-values < 0.05, p < 0.01, and p < 0.001 (*, **, and ***, respectively). Any p-values exceeding 0.05 are denoted as not significant (n.s.) or not shown.

## Acknowledgements

We thank Stephanie Ewald for her excellent technical assistance in the Core Facility Gnotobiology of the ZIEL Institute for Food & Health. We are grateful to Caroline Ziegler and Angela Sachsenhauser from the Core Facility Microbiome of the ZIEL Institute for Food & Health for outstanding technical support with 16S rRNA gene and metagenomic sequencing. We also thank Olga Seelbach and the Comparative Experimental Pathology (CEP) team for valuable discussions and technical support.

## Author contributions

D.S. and M.T. designed and supervised the study. M.T. wrote the manuscript and prepared the figures. V.B., N.B., A.E.Z., M.G.S., F. Saab, F.S., F.B., A.R., A.M., N.A.S. and M.T. interpreted and visualized data. F. Saab, F.S., F.B., A.R. and A.M. conducted bioinformatic analyses. V.B., A.E.Z., M.G.S. and M.T. performed mouse work. R.Ö. performed RNA-seq and lcWGS. K.N. performed 16S rRNA gene and metagenomic sequencing. C.M. and K.K. performed and analyzed metabolomic profiling of cecal stool samples. M.J., T.G. and K.S. performed pathological assessment. V.B., N.B., A.E.Z., M.G.S., and M.T. performed experiments. S.S., J.C.F., G.Z., D.H., R.R., and D.S. provided critical resources and input. All authors reviewed the manuscript.

## Materials availability

All unique resources generated in this study are available from the lead contact with a completed materials transfer agreement.

## Data and code availability

This paper does not report original code. All data are available from the lead contact upon request.

## Supplementary Figures

**Supplementary Figure 1.**
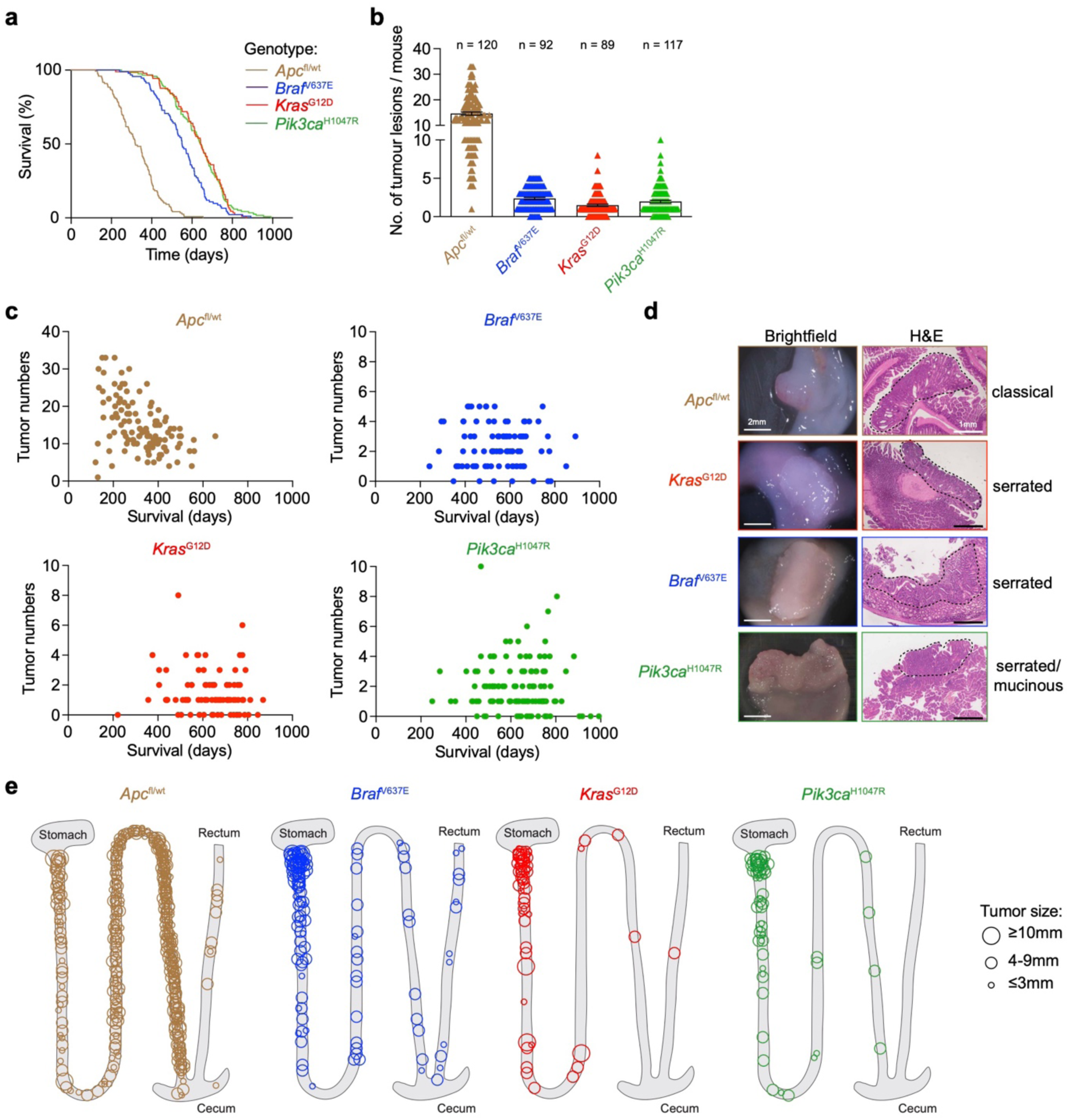
Characterization of genetically engineered intestinal cancer mouse models. **(a)** Kaplan-Meier survival analysis of *Apc*^fl/wt^, *Braf*^V637E^, *Kras*^G12D^ and *Pik3ca*^H1047R^ mice. Median survival times were 319 days for *Apc*^fl/wt^ (n = 120), 551 days for *Braf*^V637E^ (*n* = 92), 643 days for *Kras*^G12D^ (*n* = 89), and 642 days for *Pik3ca*^H1047R^ mice (*n* = 117). **(b)** Number of tumor lesions per mouse for each CRC genotype. **(c)** Tumor burden over time in *Apc*^fl/wt^, *Braf*^V637E^, *Kras*^G12D^ and *Pik3ca*^H1047R^ mice (*Apc*^fl/wt^, *n* = 120; *Braf*^V637E^, *n* = 92; *Kras*^G12D^, *n* = 89; *Pik3ca*^H1047R^, *n* = 117). **(d)** Representative macroscopic images of tumor lesions with corresponding hematoxylin and eosin (H&E)-stained sections illustrating the characteristic histopathological morphology of each CRC subtype. **(e)** Anatomical distribution of tumor lesions throughout the gastrointestinal tract for each CRC genotype. Circle diameter indicates tumor size (≤3 mm, 4-9 mm or ≥10 mm) (*Apc*^fl/wt^, *n* = 34; *Braf*^V637E^, *n* = 51; *Kras*^G12D^, *n* = 33; *Pik3ca*^H1047R^, *n* = 28).

**Supplementary Figure 2.**
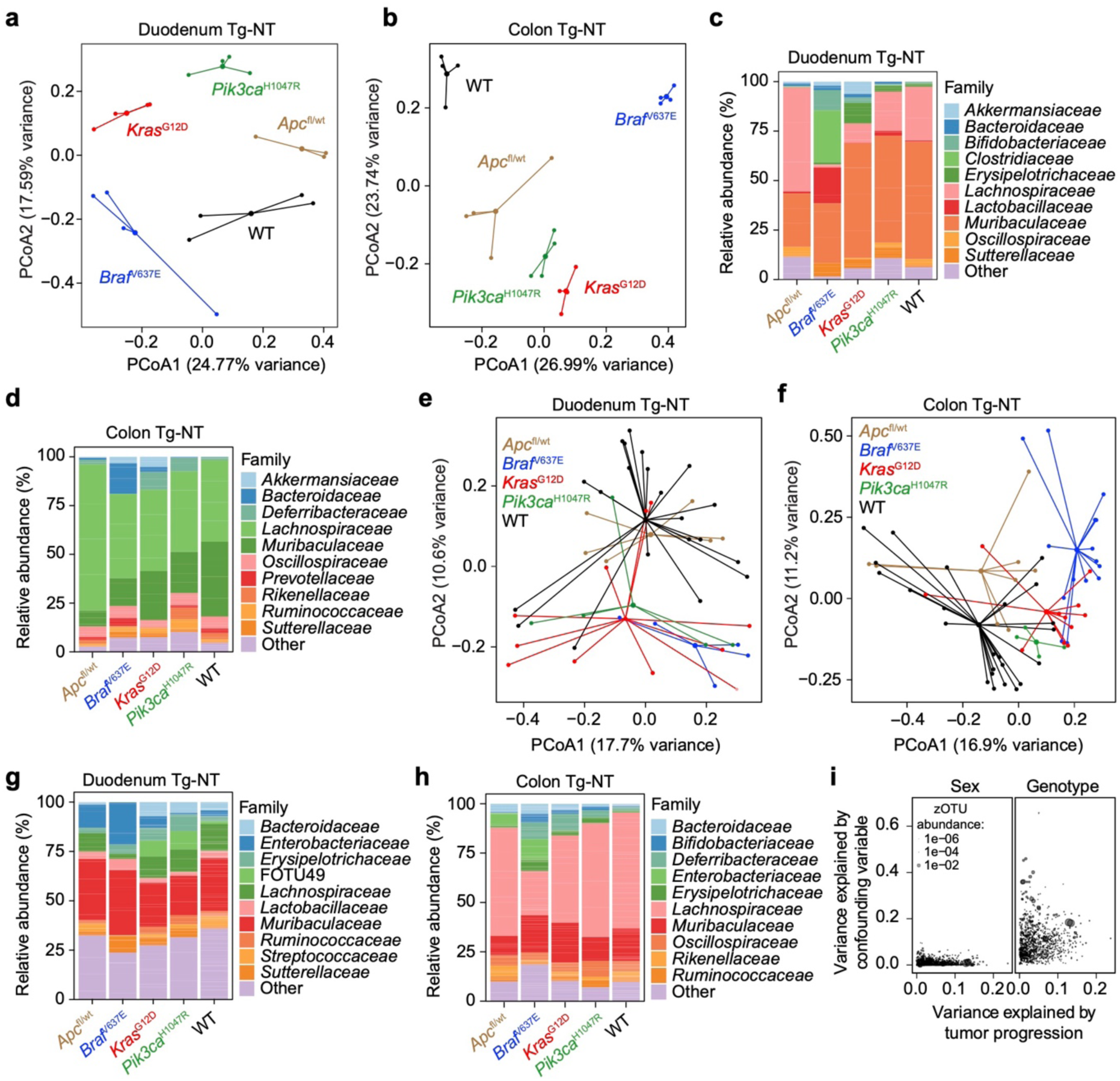
Genotype- and progression-dependent remodeling of intestinal microbial communities. **(a-b)** Principal coordinates analysis (PCoA) based on generalized UniFrac distances showing differences in beta-diversity of mucosa-associated microbial communities in the duodenum **(a)** and colon **(b)** of 8-week-old wild-type (WT) mice and non-tumorous tissue from genetically engineered intestinal cancer models (Tg-NT; *Apc*^fl/wt^, *Braf*^V637E^, *Kras*^G12D^, *Pik3ca*^H1047R^) prior to overt tumor formation (*n* = 4; PERMANOVA, p = 0.001). **(c-d)** Family-level taxonomic profiles of the mucosa-associated microbiota corresponding to the principal coordinates analyses shown in panels **a** and **b**. Relative abundances of the dominant bacterial families are shown for individual duodenal **(c)** and colonic **(d)** non-tumorous tissue (Tg-NT) from genetically engineered intestinal cancer models at 8 weeks of age. **(e-f)** Principal coordinates analysis (PCoA) based on generalized UniFrac distances showing the beta-diversity of mucosa-associated microbial communities in duodenal **(e)** and colonic **(f)** non-tumorous tissue from transgenic mice (Tg-NT) collected from tumor-bearing mice at the experimental endpoint. Lines connect samples originating from the same mouse (Duodenum/Colon; WT, *n* = 17/18; *Apc*^fl/wt^, *n* = 6/8; *Braf*^V637E^, *n* = 5/15; *Kras*^G12D^, *n* = 11/11; *Pik3ca*^H1047R^, *n* = 5/4; Duodenum, PERMANOVA, p = 0.002; Colon, PERMANOVA, p = 0.001). **(g-h)** Family-level taxonomic profiles of the mucosa-associated microbiota corresponding to the principal coordinates analyses shown in panels **e** and **f**. Relative abundances of the dominant bacterial families are shown for individual duodenal **(g)** and colonic **(h)** non-tumorous tissue samples from transgenic mice (Tg-NT) collected at the experimental endpoint (WT, *n* = 17; *Apc*^fl/wt^, *n* = 6; *Braf*^V637E^, *n* = 5; *Kras*^G12D^, *n* = 11; *Pik3ca*^H1047R^, *n* = 5). **(i)** Variance partitioning analysis of observed zero-radius operational taxonomic units (zOTUs) from small intestinal samples, showing the relative contribution of tumor progression stage (x-axis) and potential confounding variables, including sex and genotype (y-axis), to microbial community composition (*n* = 169).

**Supplementary Figure 3.**
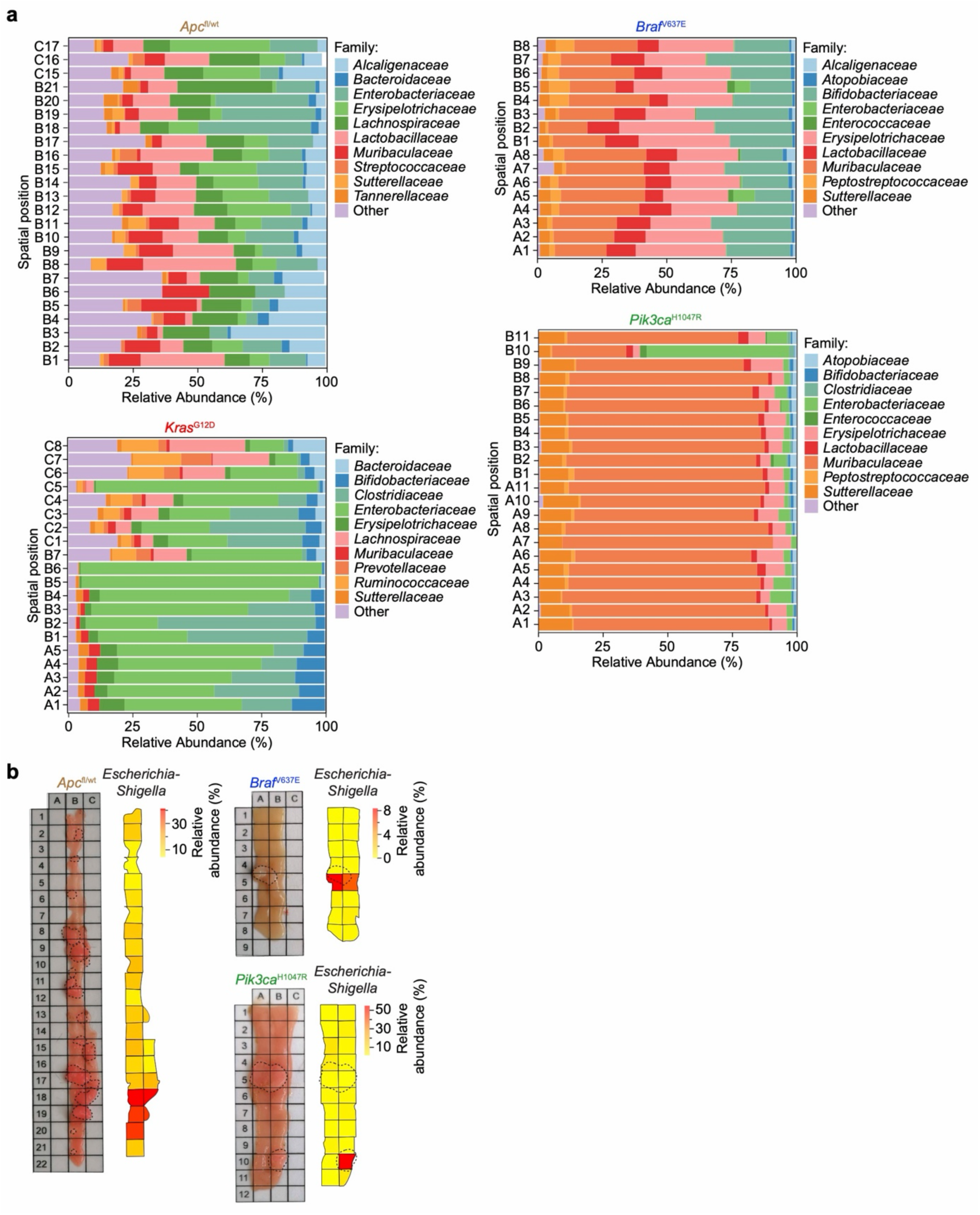
Spatially resolved 16S rRNA gene sequencing reveals tumor-associated microbial patterns in genetically engineered models of intestinal tumorigenesis. **(a)** Spatially resolved 16S rRNA gene sequencing analysis (taxonomic profiling) of the intestinal microbiota in tumor-bearing *Apc*^fl/wt^, *Braf*^V637E^, *Kras*^G12D^ and *Pik3ca*^H1047R^ mice. Horizontal bar plots show the relative abundance of bacterial families (x-axis) across individual intestinal segments (y-axis), with each segment denoted by an alphanumeric identifier (e.g., A1-A11, B1-B21, and C1-C8) corresponding to its spatial position along the dissected intestinal tract. **(b)** Representative macroscopic images of segmented intestinal samples together with the corresponding spatial distribution of *Escherichia-Shigella* inferred from 16S rRNA gene sequencing. Relative abundance values for *Escherichia-Shigella* in each intestinal segment are displayed as heatmaps, revealing distinct patterns of bacterial enrichment in tumor regions compared with adjacent non-tumorous tissue. Tumor areas are indicated by dotted lines.

**Supplementary Figure 4.**
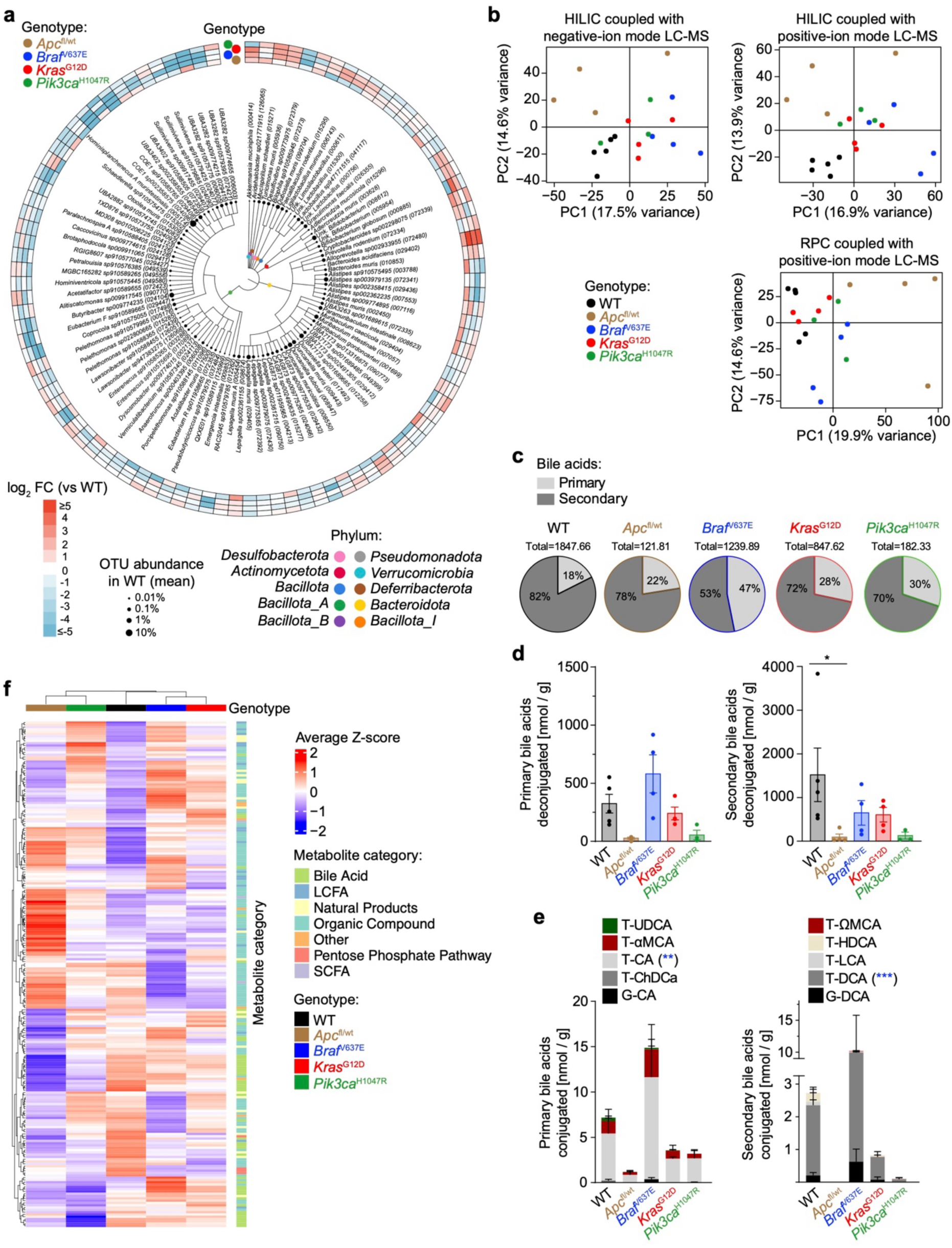
Genotype-specific metagenomic and metabolomic profiling of cecal contents across genetically engineered mouse models of intestinal tumorigenesis. **(a)** Circular cladogram of shotgun metagenomic taxonomic profiling showing bacterial taxa differentially abundant in *Apc*^fl/wt^, *Braf*^V637E^, *Kras*^G12D^ and *Pik3ca*^H1047R^ mice relative to wild-type (WT). Circle size indicates the mean OTU abundance in WT, and color intensity represents log_2_ fold change relative to WT (*n* = 3-5). **(b)** Principal component analysis (PCA) of untargeted metabolomic profiling performed using hydrophilic interaction liquid chromatography (HILIC) coupled with negative-ion liquid chromatography-mass spectrometry (LC-MS), HILIC coupled with positive-ion LC-MS, and reverse-phase chromatography (RPC) coupled with positive-ion LC-MS (*n* = 3-5). **(c)** Heatmap showing average metabolite Z-scores across CRC genotypes, with metabolites grouped according to metabolite category. **(d)** Quantification of deconjugated primary and secondary bile acids (*n* = 3-5). **(e)** Quantification of conjugated primary and secondary bile acids (*n* = 3-5; Ordinary one-way ANOVA with Bonferroni’s multiple comparisons test, * p < 0.05). **(f)** Relative contribution of primary and secondary bile acids to the total bile acid pool for each genotype (*n* = 3-5; Two-way ANOVA with Dunnett’s multiple comparisons test, ** p < 0.01, and *** p < 0.001). T-, taurine-conjugated; G-, glycine-conjugated; UDCA, ursodeoxycholic acid; αMCA, α-muricholic acid; CA, cholic acid; ChDCA, chenodeoxycholic acid; ωMCA, ω-muricholic acid; HDCA, hyodeoxycholic acid; LCA, lithocholic acid; DCA, deoxycholic acid.

**Supplementary Figure 5.**
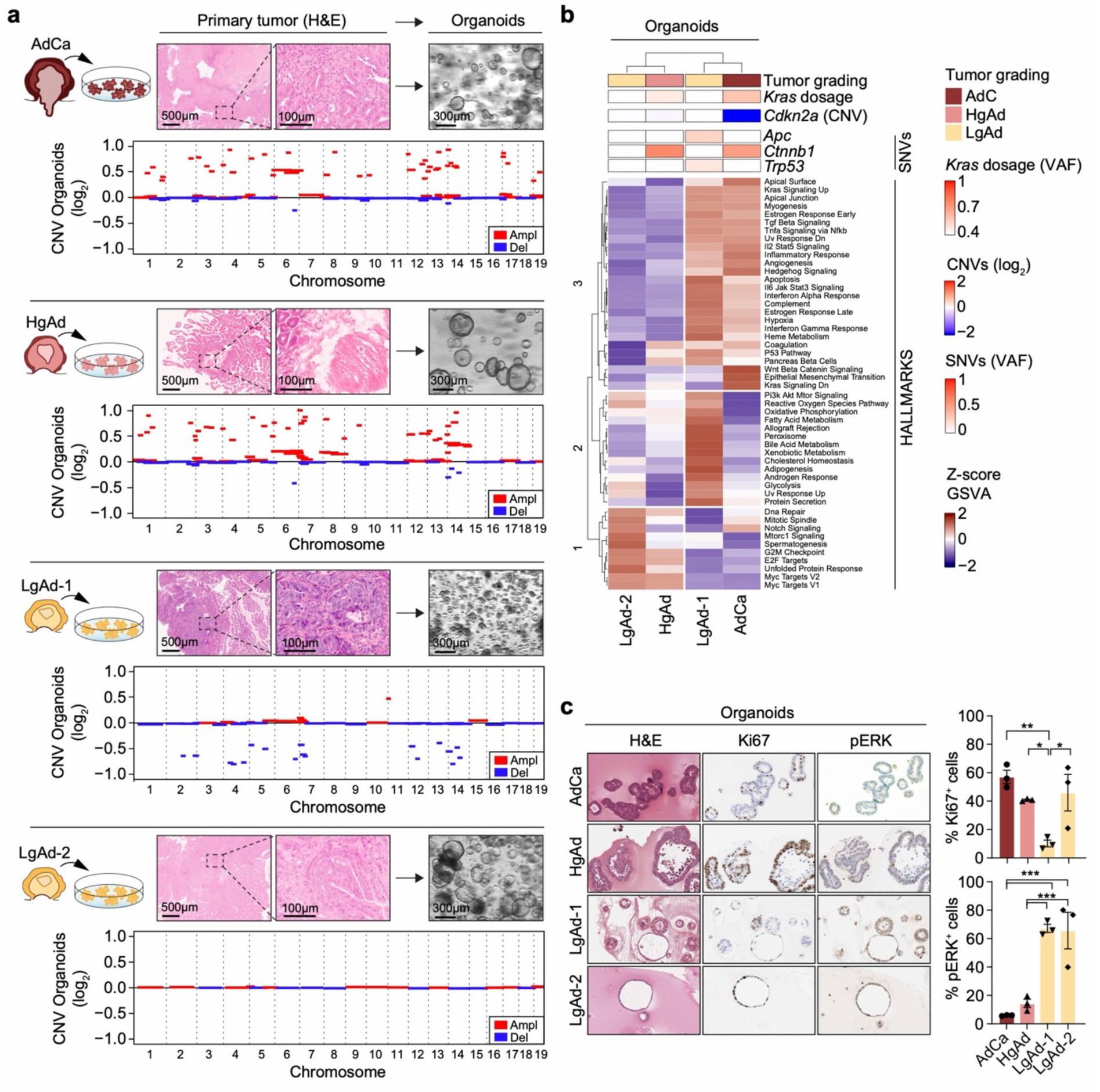
Molecular characterization of *Kras*^G12D^ tumor organoids representing distinct stages of tumor progression. **(a)** Representative workflow for the establishment of organoid cultures from adenocarcinoma (AdCa), high-grade adenoma (HgAd), low-grade adenoma 1 (LgAd-1) and low-grade adenoma 2 (LgAd-2), including representative hematoxylin and eosin (H&E)-stained sections of the primary tumors and brightfield images of the corresponding organoids. Copy number variation (CNV) profiles of the corresponding organoid lines are shown as log_2_ copy number ratios across all chromosomes, with amplifications (Ampl) and deletions (Del) indicated. **(b)** Integrated molecular characterization of organoid lines showing tumor grade, *Kras*^G12D^ variant allele frequency (VAF), *Cdkn2a* copy number variation (CNV), single nucleotide variations (SNVs) in *Apc*, *Ctnnb1* and *Trp53*, and gene set variation analysis (GSVA) of Hallmark pathway activity. **(c)** Representative hematoxylin and eosin (H&E) stainings, Ki67 and phospho-ERK (pERK) immunohistochemistry stainings of organoids together with quantification of Ki67^+^ and pERK^+^ cells (*n* = 3; Ordinary one-way ANOVA with Tukey’s multiple comparisons test, * p < 0.05, ** p < 0.01, and *** p < 0.001).

## References

1 Cancer Genome Atlas, N. Comprehensive molecular characterization of human colon and rectal cancer. Nature 487, 330–337, doi:10.1038/nature11252 (2012).

2 Linnekamp, J. F. et al. Consensus molecular subtypes of colorectal cancer are recapitulated in in vitro and in vivo models. Cell death and differentiation 25, 616–633, doi:10.1038/s41418-017-0011-5 (2018).

3 Fearon, E. R. & Vogelstein, B. A genetic model for colorectal tumorigenesis. Cell 61, 759–767, doi:10.1016/0092-8674(90)90186-i (1990).

4 Rajagopalan, H. et al. Tumorigenesis: RAF/RAS oncogenes and mismatch-repair status. Nature 418, 934, doi:10.1038/418934a (2002).

5 Chan, T. L., Zhao, W., Leung, S. Y., Yuen, S. T. & Cancer Genome, P. BRAF and KRAS mutations in colorectal hyperplastic polyps and serrated adenomas. Cancer research 63, 4878–4881 (2003).

6 Samuels, Y. et al. High frequency of mutations of the PIK3CA gene in human cancers. Science 304, 554, doi:10.1126/science.1096502 (2004).

7 Calon, A. et al. Stromal gene expression defines poor-prognosis subtypes in colorectal cancer. Nature genetics 47, 320–329, doi:10.1038/ng.3225 (2015).

8 Liao, W. et al. KRAS-IRF2 Axis Drives Immune Suppression and Immune Therapy Resistance in Colorectal Cancer. Cancer Cell 35, 559–572 e557, doi:10.1016/j.ccell.2019.02.008 (2019).

9 Lee, H. O. et al. Lineage-dependent gene expression programs influence the immune landscape of colorectal cancer. Nature genetics 52, 594–603, doi:10.1038/s41588-020-0636-z (2020).

10 Ma, J. et al. Mechanistic Foundations of KRAS-Driven Tumor Ecosystems: Integrating Crosstalk among Immune, Metabolic, Microbial, and Stromal Microenvironment. Adv Sci (Weinh) 12, e02714, doi:10.1002/advs.202502714 (2025).

11 Arthur, J. C. et al. Intestinal inflammation targets cancer-inducing activity of the microbiota. Science 338, 120–123, doi:10.1126/science.1224820 (2012).

12 Yuan, D. et al. A comprehensive analysis of the microbiota composition and host driver gene mutations in colorectal cancer. Invest New Drugs 40, 884–894, doi:10.1007/s10637-022-01263-1 (2022).

13 Kostic, A. D. et al. Genomic analysis identifies association of Fusobacterium with colorectal carcinoma. Genome research 22, 292–298, doi:10.1101/gr.126573.111 (2012).

14 Wu, S. et al. A human colonic commensal promotes colon tumorigenesis via activation of T helper type 17 T cell responses. Nature medicine 15, 1016–1022, doi:10.1038/nm.2015 (2009).

15 Rubinstein, M. R. et al. Fusobacterium nucleatum promotes colorectal carcinogenesis by modulating E-cadherin/beta-catenin signaling via its FadA adhesin. Cell host & microbe 14, 195–206, doi:10.1016/j.chom.2013.07.012 (2013).

16 Pleguezuelos-Manzano, C. et al. Mutational signature in colorectal cancer caused by genotoxic pks(+) E. coli. Nature 580, 269–273, doi:10.1038/s41586-020-2080-8 (2020).

17 Dejea, C. M. et al. Patients with familial adenomatous polyposis harbor colonic biofilms containing tumorigenic bacteria. Science 359, 592–597, doi:10.1126/science.aah3648 (2018).

18 Routy, B. et al. Gut microbiome influences efficacy of PD-1-based immunotherapy against epithelial tumors. Science 359, 91–97, doi:10.1126/science.aan3706 (2018).

19 Gopalakrishnan, V. et al. Gut microbiome modulates response to anti-PD-1 immunotherapy in melanoma patients. Science 359, 97–103, doi:10.1126/science.aan4236 (2018).

20 Shiao, S. L. et al. Commensal bacteria and fungi differentially regulate tumor responses to radiation therapy. Cancer Cell 39, 1202–1213 e1206, doi:10.1016/j.ccell.2021.07.002 (2021).

21 Zeller, G. et al. Potential of fecal microbiota for early-stage detection of colorectal cancer. Molecular systems biology 10, 766, doi:10.15252/msb.20145645 (2014).

22 Feng, Q. et al. Gut microbiome development along the colorectal adenoma-carcinoma sequence. Nature communications 6, 6528, doi:10.1038/ncomms7528 (2015).

23 Castellarin, M. et al. Fusobacterium nucleatum infection is prevalent in human colorectal carcinoma. Genome research 22, 299–306, doi:10.1101/gr.126516.111 (2012).

24 Yachida, S. et al. Metagenomic and metabolomic analyses reveal distinct stage-specific phenotypes of the gut microbiota in colorectal cancer. Nature medicine 25, 968–976, doi:10.1038/s41591-019-0458-7 (2019).

25 Wirbel, J. et al. Meta-analysis of fecal metagenomes reveals global microbial signatures that are specific for colorectal cancer. Nature medicine 25, 679–689, doi:10.1038/s41591-019-0406-6 (2019).

26 Thomas, A. M., et al. Metagenomic analysis of colorectal cancer datasets identifies cross-cohort microbial diagnostic signatures and a link with choline degradation. Nature medicine 25, 667–678, doi:10.1038/s41591-019-0405-7 (2019).

27 Taketo, M. M. & Edelmann, W. Mouse models of colon cancer. Gastroenterology 136, 780–798, doi:10.1053/j.gastro.2008.12.049 (2009).

28 Dienstmann, R. et al. Consensus molecular subtypes and the evolution of precision medicine in colorectal cancer. Nature reviews. Cancer 17, 79–92, doi:10.1038/nrc.2016.126 (2017).

29 Guinney, J. et al. The consensus molecular subtypes of colorectal cancer. Nature medicine 21, 1350–1356, doi:10.1038/nm.3967 (2015).

30 Vogelstein, B. et al. Genetic alterations during colorectal-tumor development. The New England journal of medicine 319, 525–532, doi:10.1056/NEJM198809013190901 (1988).

31 Bennecke, M. et al. Ink4a/Arf and oncogene-induced senescence prevent tumor progression during alternative colorectal tumorigenesis. Cancer Cell 18, 135–146, doi:10.1016/j.ccr.2010.06.013 (2010).

32 Rad, R. et al. A genetic progression model of Braf(V600E)-induced intestinal tumorigenesis reveals targets for therapeutic intervention. Cancer Cell 24, 15–29, doi:10.1016/j.ccr.2013.05.014 (2013).

33 Itzkowitz, S. H. & Yio, X. Inflammation and cancer IV. Colorectal cancer in inflammatory bowel disease: the role of inflammation. American journal of physiology. Gastrointestinal and liver physiology 287, G7–17, doi:10.1152/ajpgi.00079.2004 (2004).

34 Eser, S. et al. Selective requirement of PI3K/PDK1 signaling for Kras oncogene-driven pancreatic cell plasticity and cancer. Cancer Cell 23, 406–420, doi:10.1016/j.ccr.2013.01.023 (2013).

35 Cheung, A. F. et al. Complete deletion of Apc results in severe polyposis in mice. Oncogene 29, 1857–1864, doi:10.1038/onc.2009.457 (2010).

36 O’Keefe, S. J. Diet, microorganisms and their metabolites, and colon cancer. Nature reviews. Gastroenterology & hepatology 13, 691–706, doi:10.1038/nrgastro.2016.165 (2016).

37 Roper, J. et al. Colonoscopy-based colorectal cancer modeling in mice with CRISPR-Cas9 genome editing and organoid transplantation. Nature protocols 13, 217–234, doi:10.1038/nprot.2017.136 (2018).

38 Atatri, S. et al. Protocol for sub-mucosal orthotopic injection of organoids into murine colon to study tumor growth and metastasis. STAR Protoc 6, 103887, doi:10.1016/j.xpro.2025.103887 (2025).

39 Fumagalli, A. et al. Genetic dissection of colorectal cancer progression by orthotopic transplantation of engineered cancer organoids. Proceedings of the National Academy of Sciences of the United States of America 114, E2357–E2364, doi:10.1073/pnas.1701219114 (2017).

40 Nicolas, A. M. et al. Inflammatory fibroblasts mediate resistance to neoadjuvant therapy in rectal cancer. Cancer Cell 40, 168–184 e113, doi:10.1016/j.ccell.2022.01.004 (2022).

41 Richon, S., Zajac, O., Perez Gonzalez, C. & Matic Vignjevic, D. Optimized protocol for the generation of an orthotopic colon cancer mouse model and metastasis. STAR Protoc 4, 102022, doi:10.1016/j.xpro.2022.102022 (2023).

42 Watanabe, S. et al. Transplantation of intestinal organoids into a mouse model of colitis. Nature protocols 17, 649–671, doi:10.1038/s41596-021-00658-3 (2022).

43 O’Rourke, K. P. et al. Transplantation of engineered organoids enables rapid generation of metastatic mouse models of colorectal cancer. Nature biotechnology 35, 577–582, doi:10.1038/nbt.3837 (2017).

44 Felchle, H. et al. Novel Tumor Organoid-Based Mouse Model to Study Image Guided Radiation Therapy of Rectal Cancer After Noninvasive and Precise Endoscopic Implantation. International journal of radiation oncology, biology, physics 118, 1094–1104, doi:10.1016/j.ijrobp.2023.10.008 (2024).

45 Mueller, S. et al. A disease model resource reveals core principles of tissue-specific cancer evolution. Nature 653, 265–276, doi:10.1038/s41586-026-10187-2 (2026).

46 Mazmanian, S. K., Liu, C. H., Tzianabos, A. O. & Kasper, D. L. An immunomodulatory molecule of symbiotic bacteria directs maturation of the host immune system. Cell 122, 107–118, doi:10.1016/j.cell.2005.05.007 (2005).

47 Ivanov, II et al. Specific microbiota direct the differentiation of IL-17-producing T-helper cells in the mucosa of the small intestine. Cell host & microbe 4, 337–349, doi:10.1016/j.chom.2008.09.009 (2008).

48 Atarashi, K. et al. Induction of colonic regulatory T cells by indigenous Clostridium species. Science 331, 337–341, doi:10.1126/science.1198469 (2011).

49 Belcheva, A. et al. Gut microbial metabolism drives transformation of MSH2-deficient colon epithelial cells. Cell 158, 288–299, doi:10.1016/j.cell.2014.04.051 (2014).

50 Pekel, S. et al. Meta-analysis reveals microbiome signatures for colorectal cancer that are universal across age groups and sequencing methods. Cell host & microbe, doi:10.1016/j.chom.2026.05.030 (2026).

51 Vaishnava, S. et al. The antibacterial lectin RegIIIgamma promotes the spatial segregation of microbiota and host in the intestine. Science 334, 255–258, doi:10.1126/science.1209791 (2011).

52 Salzman, N. H. et al. Enteric defensins are essential regulators of intestinal microbial ecology. Nature immunology 11, 76–83, doi:10.1038/ni.1825 (2010).

53 Fujii, M. et al. Human Intestinal Organoids Maintain Self-Renewal Capacity and Cellular Diversity in Niche-Inspired Culture Condition. Cell stem cell 23, 787–793 e786, doi:10.1016/j.stem.2018.11.016 (2018).

54 Drost, J. et al. Sequential cancer mutations in cultured human intestinal stem cells. Nature 521, 43–47, doi:10.1038/nature14415 (2015).

55 Matano, M. et al. Modeling colorectal cancer using CRISPR-Cas9-mediated engineering of human intestinal organoids. Nature medicine 21, 256–262, doi:10.1038/nm.3802 (2015).

56 Bouskra, D. et al. Lymphoid tissue genesis induced by commensals through NOD1 regulates intestinal homeostasis. Nature 456, 507–510, doi:10.1038/nature07450 (2008).

57 Yu, T. et al. Fusobacterium nucleatum Promotes Chemoresistance to Colorectal Cancer by Modulating Autophagy. Cell 170, 548–563 e516, doi:10.1016/j.cell.2017.07.008 (2017).

58 Mima, K. et al. Fusobacterium nucleatum in colorectal carcinoma tissue and patient prognosis. Gut 65, 1973–1980, doi:10.1136/gutjnl-2015-310101 (2016).

59 Madison, B. B. et al. Cis elements of the villin gene control expression in restricted domains of the vertical (crypt) and horizontal (duodenum, cecum) axes of the intestine. The Journal of biological chemistry 277, 33275–33283, doi:10.1074/jbc.M204935200 (2002).

60 Nagtegaal, I. D. et al. The 2019 WHO classification of tumours of the digestive system. Histopathology 76, 182–188, doi:10.1111/his.13975 (2020).

61 Jesinghaus, M. et al. A translational colorectal cancer organoid biobank mirrors patients’ tumor histology, molecular profiles, and treatment responses. J Exp Clin Cancer Res 45, doi:10.1186/s13046-026-03666-x (2026).

62 Parekh, S., Ziegenhain, C., Vieth, B., Enard, W. & Hellmann, I. The impact of amplification on differential expression analyses by RNA-seq. Scientific reports 6, 25533, doi:10.1038/srep25533 (2016).

63 Jesinghaus, M. et al. Morphology Matters: A Critical Reappraisal of the Clinical Relevance of Morphologic Criteria From the 2019 WHO Classification in a Large Colorectal Cancer Cohort Comprising 1004 Cases. Am J Surg Pathol 45, 969–978, doi:10.1097/PAS.0000000000001692 (2021).

64 Reitmeier, S., Kiessling, S., Neuhaus, K. & Haller, D. Comparing Circadian Rhythmicity in the Human Gut Microbiome. STAR Protoc 1, 100148, doi:10.1016/j.xpro.2020.100148 (2020).

65 Lagkouvardos, I. et al. IMNGS: A comprehensive open resource of processed 16S rRNA microbial profiles for ecology and diversity studies. Scientific reports 6, 33721, doi:10.1038/srep33721 (2016).

66 Kioukis, A., Pourjam, M., Neuhaus, K. & Lagkouvardos, I. Taxonomy Informed Clustering, an Optimized Method for Purer and More Informative Clusters in Diversity Analysis and Microbiome Profiling. Front Bioinform 2, 864597, doi:10.3389/fbinf.2022.864597 (2022).

67 Dietrich, A. et al. Namco: a microbiome explorer. Microb Genom 8, doi:10.1099/mgen.0.000852 (2022).

68 Lagkouvardos, I., Fischer, S., Kumar, N. & Clavel, T. Rhea: a transparent and modular R pipeline for microbial profiling based on 16S rRNA gene amplicons. PeerJ 5, e2836, doi:10.7717/peerj.2836 (2017).

69 Zhang, J. et al. spatialHeatmap: visualizing spatial bulk and single-cell assays in anatomical images. NAR Genom Bioinform 6, lqae006, doi:10.1093/nargab/lqae006 (2024).

70 Ruscheweyh, H. J. et al. Cultivation-independent genomes greatly expand taxonomic-profiling capabilities of mOTUs across various environments. Microbiome 10, 212, doi:10.1186/s40168-022-01410-z (2022).

71 Mallick, H. et al. Multivariable association discovery in population-scale meta-omics studies. PLoS Comput Biol 17, e1009442, doi:10.1371/journal.pcbi.1009442 (2021).

72 Letunic, I. & Bork, P. Interactive Tree of Life (iTOL) v6: recent updates to the phylogenetic tree display and annotation tool. Nucleic acids research 52, W78–W82, doi:10.1093/nar/gkae268 (2024).

73 Kessner, D., Chambers, M., Burke, R., Agus, D. & Mallick, P. ProteoWizard: open source software for rapid proteomics tools development. Bioinformatics 24, 2534–2536, doi:10.1093/bioinformatics/btn323 (2008).

74 Smith, C. A., Want, E. J., O’Maille, G., Abagyan, R. & Siuzdak, G. XCMS: processing mass spectrometry data for metabolite profiling using nonlinear peak alignment, matching, and identification. Analytical chemistry 78, 779–787, doi:10.1021/ac051437y (2006).

75 Tsugawa, H. et al. MS-DIAL: data-independent MS/MS deconvolution for comprehensive metabolome analysis. Nature methods 12, 523–526, doi:10.1038/nmeth.3393 (2015).

76 Gu, Z. Complex heatmap visualization. Imeta 1, e43, doi:10.1002/imt2.43 (2022).

77 Reiter, S. et al. Development of a Highly Sensitive Ultra-High-Performance Liquid Chromatography Coupled to Electrospray Ionization Tandem Mass Spectrometry Quantitation Method for Fecal Bile Acids and Application on Crohn’s Disease Studies. J Agric Food Chem 69, 5238–5251, doi:10.1021/acs.jafc.1c00769 (2021).

78 Han, J., Lin, K., Sequeira, C. & Borchers, C. H. An isotope-labeled chemical derivatization method for the quantitation of short-chain fatty acids in human feces by liquid chromatography-tandem mass spectrometry. Anal Chim Acta 854, 86–94, doi:10.1016/j.aca.2014.11.015 (2015).

